# Mapping grey and white matter activity in the human brain with isotropic ADC-fMRI

**DOI:** 10.1101/2024.10.01.615823

**Authors:** Arthur P C Spencer, Jasmine Nguyen-Duc, Inès de Riedmatten, Filip Szczepankiewicz, Ileana O Jelescu

## Abstract

Functional MRI (fMRI) using the blood-oxygen level dependent (BOLD) signal provides valuable insight into grey matter activity. However, uncertainty surrounds the white matter BOLD signal. Apparent diffusion coefficient (ADC) offers an alternative fMRI contrast sensitive to transient cellular deformations during neural activity, facilitating detection of both grey and white matter activity. Further, through minimising vascular contamination, ADC-fMRI has the potential to overcome the limited temporal specificity of the BOLD signal. However, the use of linear diffusion encoding introduces sensitivity to fibre directionality, while averaging over multiple directions comes at great cost to temporal resolution. In this study, we used spherical b-tensor encoding to impart diffusion sensitisation in all directions per shot, providing an ADC-fMRI contrast capable of detecting activity independently of fibre directionality. We provide evidence from two task-based experiments on a clinical scanner that isotropic ADC-fMRI is more temporally specific than BOLD-fMRI, and offers more balanced mapping of grey and white matter activity. We further demonstrate that isotropic ADC-fMRI detects white matter activity independently of fibre direction, while linear ADC-fMRI preferentially detects activity in voxels containing fibres perpendicular to the diffusion encoding direction. Thus, isotropic ADC-fMRI opens avenues for investigation into whole-brain grey and white matter functional connectivity.

## Introduction

Non-invasive and direct detection of neural activity in vivo remains a significant challenge in neuroimaging. While functional MRI (fMRI) has substantially advanced our understanding of brain function, it is most commonly acquired using gradient-echo sequences yielding blood-oxygen level dependent (BOLD) contrast via *T* ^⇤^ weighting [1].= This carries a number of inherent limitations due to its reliance on neurovascular coupling, including limited spatial and temporal specificity [2, 3]. Additionally, due to the reduced vasculature, different energy requirements, and altered haemodynamic response in white matter, previous fMRI studies commonly attributed the white matter BOLD signal to noise, treating it as a nuisance regressor [4]. There is emerging evidence of neural activity being represented in the white matter BOLD signal in both resting state and task fMRI, however most studies of white matter BOLD signal have tailored analysis methods specifically to white matter activation, obscuring simultaneous detection of both grey and white matter activity [5–7] (note that here we refer to “neural activity” in white matter, meaning the propagation of action potentials for the transfer of information between grey matter regions carrying out neural processing). Other functional neuroimaging methods also have limited scope for detection of white matter function. For example, the sensitivity of electro- and magneto-encephalography decays rapidly with distance from the scalp, precluding accurate mapping of white matter activity [8]. Since white matter function holds considerable diagnostic value for neurological and psychiatric diseases [9], the inability to reliably map grey and white matter simultaneously represents a substantial limitation of current non-invasive functional methods.

Diffusion-weighted functional MRI (dfMRI) has the potential to overcome these limitations by detecting alterations to the diffusion of water molecules caused by morphological changes occurring during neural activity [10, 11]. These sub-micrometer scale neuromorphological changes to neuronal cell bodies [12, 13], neurites [13, 14], synaptic boutons [14], astrocytic processes [15] and axons [16] occur on a timescale of milliseconds but can last several seconds upon sustained firing, as observed with optical microscopy. Measuring these changes with dfMRI allows detection of neural activity in vivo with better temporal specificity than BOLD-fMRI, evidenced by an earlier response and faster return to baseline in response to task stimuli [17–19].

The diffusion-sensitised spin-echo MRI signal with echo time *T_E_* is characterised in the Gaussian phase approximation by

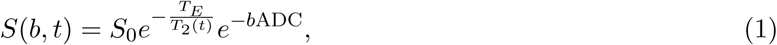

therefore a conventional dfMRI acquisition is sensitive to both apparent diffusion coefficient (ADC) and *T*_2_ changes, capturing both neuromorphological and neurovascular (BOLD) fluctuations. Although several previous studies have used dfMRI acquisitions with high b-values to increase sensitivity to ADC changes, it has been demonstrated that this signal remains sensitive to BOLD effects [20]. More recently, ADC has been used as a functional contrast in order to further eliminate vascular contributions to the diffusion signal, increasing specificity of the contrast to neural activity [21–25].

ADC-fMRI aims to eliminate the influence of the vasculature on the functional contrast via the *T*_2_ weighting, by taking the ratio of two volumes acquired at different b-values and assuming *T*_2_(*t*+*<t*) ⇡ *T*_2_(*t*), giving

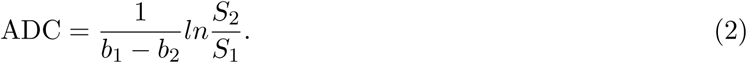

An additional source of vascular contributions to this signal can arise from the blood water pool in each voxel, where the increase in blood volume and flow during the haemodynamic response can translate into a change in ADC [21, 22, 25]. However, this contribution can be minimised by choosing both b-values 2’:200 s mm^-2^, thus largely suppressing the blood water signal in all the diffusion-weighted images acquired [26]. This approach of avoiding the perfusion regime in the ADC calculation has previously been described as ‘synthetic’ or ‘shifted’ ADC [27, 28]. With this shifted ADC approach, an ADC-fMRI acquisition can be designed to directly measures changes in the intra- and extra-cellular water diffusion, which arise due to neuromorphological changes, whilst minimising contamination with signals from vascular sources [22, 23]. Due to its sensitivity to neuromorphological coupling, ADC-fMRI is theoretically capable of detecting both grey and white matter activity; in a preclinical experiment, an ADC drop was reported in the mouse optic nerve during visual stimulation only when the linear diffusion encoding gradient direction was aligned perpendicular to optic nerve fibres, and not when it was aligned parallel to them [29, 30]. The temporal resolution of the experiment (12.8 minutes) only enabled the estimation of one ADC value pre-, during and post-stimulus, precluding the analysis of a temporal response and possible habituation effects. Linear ADC-fMRI acquisitions have since been introduced on clinical scanners with reasonable temporal resolution for fMRI [24, 25]. However, as shown in Spees et al. [29], with linear diffusion encoding, sensitivity to activity within voxels containing organised white matter fibres depends on the angle between the diffusion encoding direction and the fibre direction, requiring acquisition of multiple diffusion directions to achieve directional independence [21, 22, 31]. In this study, we use spherical b-tensor encoding [32–35] to sensitise the signal to diffusivity in all directions for every signal acquisition. With this isotropic ADC-fMRI acquisition, we achieve uniform sensitivity to activity-driven microstructural fluctuations in grey and white matter, as well as sufficient spatial and temporal resolution to enable a voxel-wise general linear model analysis of ADC-fMRI timecourses in the human brain. In particular, we aimed to detect activity throughout the optic radiation and primary visual cortex in response to visual stimulation, and throughout the corticospinal tract and the hand portion of the primary motor cortex in response to a motor task. We show that ADC-fMRI provides better temporal specificity than BOLD-fMRI whilst overcoming the dependence on white matter fibre direction exhibited by linear ADC-fMRI [36]. Thus, we provide an isotropic ADC-fMRI contrast sensitive to activity-related neuromophological changes in grey and white matter, independent of tissue organisation and with substantially improved temporal resolution compared to averaging multiple linear diffusion encoding directions.

## Results

### Visual Stimulation Task

We acquired isotropic ADC-fMRI data with alternating b-values of 200 and 1000 s mm^-2^, with spherical b-tensor encoding [34], during a flashing checkerboard visual stimulation task (n = 12 subjects). For comparison, in a subset of subjects, we also acquired linear ADC-fMRI (n = 10) using a twice-refocused spin-echo EPI sequence with bipolar linear encoding gradients, and multi-echo gradient echo BOLD-fMRI (n = 7). Detailed acquisition parameters are given in the Methods. For both linear encoding and isotropic encoding, we analysed the ADC-fMRI timeseries calculated using Equation 2 in addition to analysing each b-value timeseries (b200-dfMRI and b1000-dfMRI) which are sensitive to both ADC changes and *T*_2_ BOLD effects. Using a general linear model with the task modelled as a boxcar function, we investigated task-associated ADC decreases in ADC-fMRI and signal increases in b200-dfMRI, b1000- dfMRI and BOLD-fMRI. Spatial maps showing group-level activation during the visual stimulation task are shown in Figure 1 and in Figures S1-S4. Subject-specific spatial maps are shown in Figures S5 and S6. For completeness, BOLD-fMRI group-level spatial maps are shown when convolving the task timeseries with the haemodynamic response function (Figure S1), and ADC-fMRI group-level spatial maps are shown when convolving the task timeseries with the diffusion response function [17] (Figure S2).

**Figure 1:**
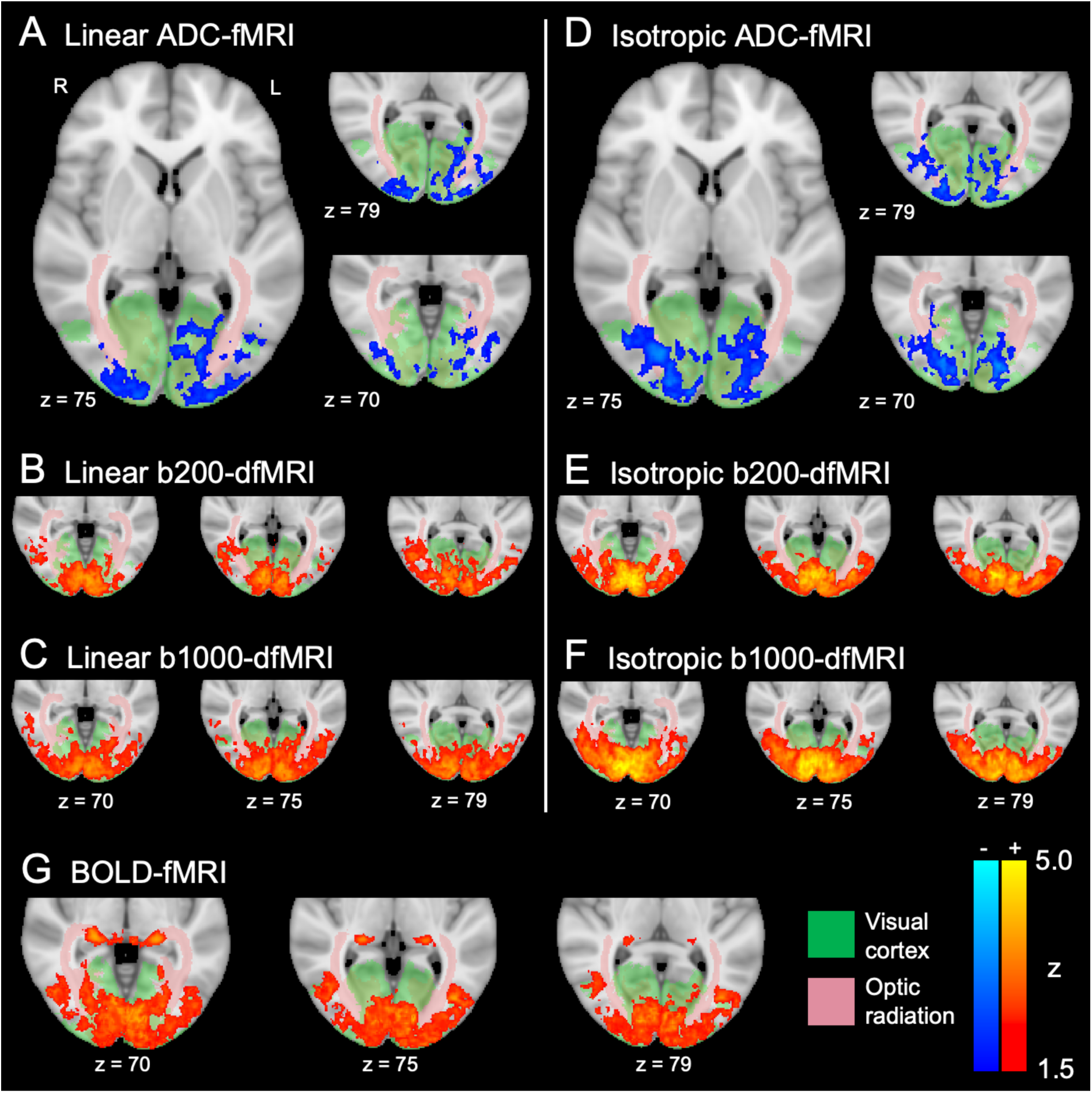
Visual task spatial maps. Colour bars show z-scores following group-level cluster correction (*z* 2’: 1.5, p *<* 0.05) for ADC-fMRI, b200-dfMRI and b1000-dfMRI with linear encoding (A, B and C respectively; n = 10) and isotropic encoding (D, E and F respectively; n = 12), in addition to BOLD- fMRI (G; n = 7). For anatomical reference, Juelich atlas regions defining the visual cortex and optic radiation are overlaid on the MNI152 standard template.

BOLD-fMRI spatial activation is localised to the visual cortex and lateral geniculate nucleus, with few active voxels in the optic radiation. Conversely, ADC-fMRI spatial maps for both linear and isotropic encoding show activation clusters in the optic radiation as well as grey matter in the visual cortex. However, activation was not detected along the entire length of the optic radiation, likely due to signal-to-noise ratio (SNR) limitations (see SNR maps in Figures S17 and S18). While the temporal SNR of BOLD-fMRI was *>*60 throughout the optic radiation, that of isotropic ADC-fMRI ranged from 30 close to the cortex to 15 near the thalamus, and was even lower for linear ADC-fMRI.

The isotropic ADC-fMRI spatial map is more symmetrical than linear ADC-fMRI, reflecting the independence of fibre direction of the isotropic ADC-fMRI sequence. For both linear and isotropic encoding, the b200-dfMRI spatial maps resemble those of BOLD-fMRI, with the majority of active voxels in the visual cortex, while b1000-dfMRI captures a combination of the diffusivity changes seen in the ADC-fMRI maps and the changes seen in the b200-dfMRI spatial maps due to the *T*_2_ weighting of the dfMRI signal, thus displaying both grey and white matter activation.

To compare sensitivity to activity in grey and white matter, we measured the proportion of significant voxels from subject-level cluster-corrected spatial maps (Figure S5 and S6) in grey or white matter, determined from the T1 image of each subject. In BOLD-fMRI spatial maps, 12.4% of significant voxels were in white matter. In linear ADC-fMRI spatial maps, 43.0% of significant voxels were in white matter, compared to 23.5% with linear b200-dfMRI and 23.6% with linear b1000-dfMRI. In isotropic ADC-fMRI spatial maps, 46.0% of significant voxels were in white matter, compared to 17.5% with isotropic b200- dfMRI and 20.6% with isotropic b1000-dfMRI. Statistical comparison at the subject level showed that ADC-fMRI detected a significantly higher proportion of white matter voxels than BOLD-fMRI, b200-dfMRI, and b1000-dfMRI, for both linear and isotropic encoding (p *<* 0.05; Figure S7, Table S3).

In a supplementary experiment following methods proposed in Schilling et al. [37], we found that the power spectrum of the mean BOLD-fMRI timecourse within the optic radiation contained significant power at the task frequency (see Supplementary Materials “Model-Free Frequency Analysis”). Eroding the region of interest to only include the core of the optic radiation revealed an altered BOLD response in deep white matter. Despite finding significant power at the task frequency in a region of interest, the optic radiation was not highlighted by voxel-wise general linear model analysis of BOLD-fMRI timeseries. Figure 2 shows the average ADC and signal responses to the visual task. Peak response amplitudes are shown in Table S1, and the time to reach 50% activation from task onset or offset are shown in Table S2. The time from task onset to 50% peak activation was quicker for both linear ADC-fMRI (1.3 s on the average response) and isotropic ADC-fMRI (1.5 s) than BOLD-fMRI (4.1 s). The rise time of b200-dfMRI (1.4 s linear, 2.6 s isotropic) and b1000-dfMRI (1.6 s linear, 2.1 s isotropic) was also quicker than BOLD-fMRI. However, both linear and isotropic b200-dfMRI and b1000-dfMRI had a slow decline from task offset to 50% activation (4.2-4.7 s) approaching that of BOLD-fMRI (5.7 s). The return to 50% activation was slightly slower for isotropic ADC-fMRI (1.8 s) than linear ADC-fMRI (1.1 s), possibly reflecting some vascular contamination of the signal in the former due to the lack of compensation for cross-terms between diffusion-weighting gradients and susceptibility-induced background gradient fields (see Discussion). Comparison of rise and fall times at the subject level showed that these differences between contrasts were significant (p *<* 0.05; Figure S8, Table S4). When measuring the response averaged over significant voxels within subject-specific grey and white matter tissues maps, the amplitude of the average BOLD response was lower in white matter (1.2%; Figure 2I) than grey matter (2.7%; Figure 2F), reflecting the differences between the vascularisation of each tissue. The amplitude of the average linear ADC-fMRI response was similar in grey and white matter (-1.3%; Figures 2D and 2G). With isotropic encoding, this response was only slightly smaller in white matter (-1.0%; Figure 2H) compared to grey matter (-1.2%; Figure 2E). In white matter, the average response with isotropic ADC-fMRI was slightly lower in amplitude than linear ADC-fMRI (-1.0% vs -1.3%). This could be due to isotropic encoding measuring average ADC changes over all spatial directions (therefore including the smaller changes in diffusion parallel to fibres), in contrast with linear encoding measurements which may be significant only when maximally sensitive to fibres perpendicular to the diffusion encoding direction.

**Figure 2:**
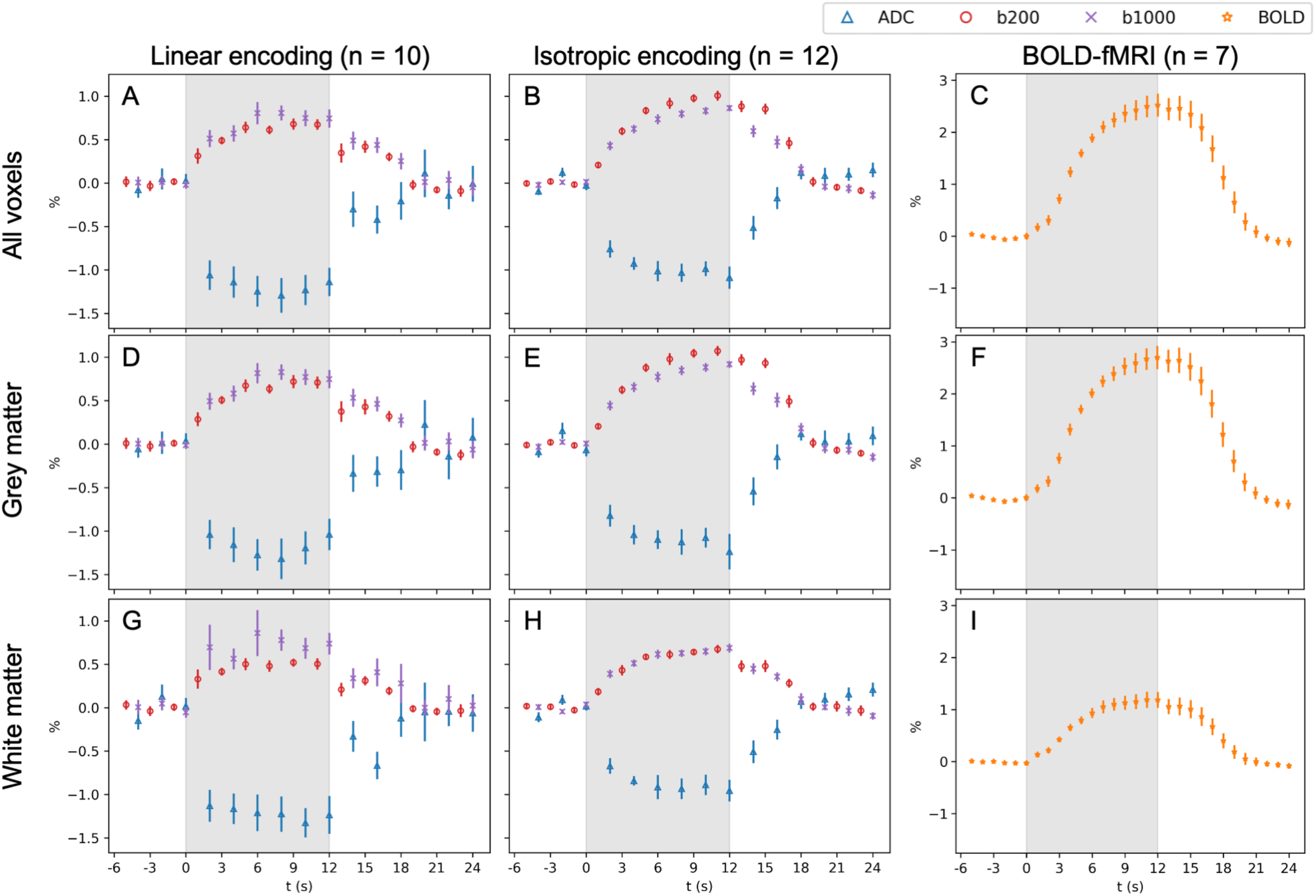
Visual task response in active voxels from subject-level spatial maps. Plots show the average timecourse, with bars indicating the standard error of the mean across subjects (linear n = 10; isotropic n = 12; BOLD n = 7). Subject-specific timecourses were averaged across epochs and across voxels in spatial maps following subject-level cluster correction (*z* 2’: 2.3, p *<* 0.05). Time is given in reference to the task onset, with the task stimulation duration indicated by the shaded area. This is shown for all significant voxels (A-C) and for significant voxels within subject-specific maps of grey (D-F) and white matter (G-I).

### Directionality

We then aimed to determine the sensitivity of ADC-fMRI acquisitions to white matter fibre organisation. To measure the directionality of white matter fibres, we acquired multi-shell diffusion-weighted imaging data in each subject and performed constrained spherical deconvolution to obtain the fibre orientation distribution (FOD) in each voxel (see Methods). We defined the fibre direction as the direction of the largest FOD peak, then measured the angle between the fibre direction and the diffusion encoding direction from the linear ADC-fMRI acquisition (which was also used as the reference direction for the spherical b-tensor encoding waveform).

In voxels active during visual stimulation, the distribution of fibre angles differed between isotropic and linear ADC-fMRI (p *<* 0.0001, two-sample Kolmogorov-Smirnov test), with linear ADC-fMRI displaying a preference for detecting activity in voxels containing fibres more perpendicular to the diffusion encoding direction (Figure 3A). Additionally, while the percentage ADC change measured in each significant voxel with isotropic ADC-fMRI was consistent across fibre angles, linear ADC-fMRI yielded larger magnitude ADC changes at higher angles (Figure 3B). This exemplifies the increased sensitivity of linear ADC-fMRI in voxels containing fibres perpendicular to the diffusion encoding direction.

**Figure 3:**
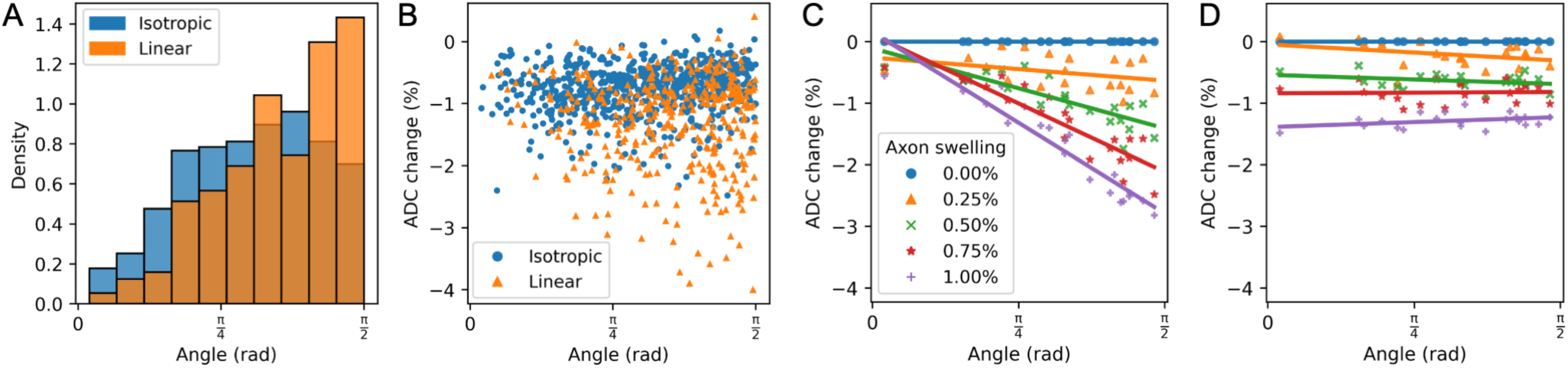
Comparison of the dependence of linear vs isotropic ADC-fMRI on white matter fibre angle. The diffusion encoding direction from the linear dfMRI sequence was used as a reference to measure the angle to the largest FOD peak in each active voxel in subject-level cluster-corrected spatial maps (*z* 2’: 2.3, p *<* 0.05), shown as a density distribution of significant voxels as a function of angle (A) and as the percentage change in ADC during visual stimulation for each voxel, as a function of angle (B). Simulated ADC changes when in silico axons swell to 0-1% of their original volume are plotted as a function of angle for linear ADC-fMRI (C) and isotropic ADC-fMRI (D).

### *In Silico* Experiments

To assess the effect of fibre angle on the ADC changes measured by linear and isotropic ADC-fMRI, we carried out in silico diffusion MRI experiments using the Monte Carlo Diffusion Simulator [38] on numerical white matter phantoms of densely-packed realistic axons generated by the CATERPillar tool [39]. Axons were swollen (by 0.25%, 0.5%, 0.75%, and 1% of their original volume) and ADC was measured as a percentage change from baseline. In linear ADC-fMRI simulations using a pulsed gradient pair, there was a strong dependence of ADC change on the angle between the fibre population and the diffusion encoding direction (Figure 3C), with the trend becoming steeper with larger amounts of swelling. In isotropic ADC-fMRI simulations, the ADC change was consistent across fibre angles (Figure 3D). Figure S10 shows the simulated intracellular and extracellular ADC changes as a function of axonal volume swelling and angle, separately for each compartment, which showcases that most of the “combined” ADC change is driven by a change in the extracellular water with swelling, the latter also showing a more marked dependence on fibre angle. Figure S11 shows the intracellular, extracellular and combined baseline ADC values (i.e. with no cell swelling) measured at different fibre angles, and the intracellular volume fraction at each swelling percentage. This demonstrates that the combined ADC drop is also, to a smaller extent, influenced by an increase in the intracellular water fraction (which has comparably lower diffusivity than the extracellular water). Additionally, this shows that the measured ADC difference between intracellular and extracellular water is greater at higher linear encoding angles to the fibre. Therefore, a given increase in intracellular volume fraction would result in a larger ADC change at a higher angle to the fibre. Thus, both of these mechanisms whereby the cell swelling influences the measured ADC change (the changes to extracellular diffusivity, and the increase in intracellular contributions to the signal) are dependent on fibre angle.

### Motor Task

To confirm that these results are not specific to the visual system, we also acquired isotropic ADC-fMRI and BOLD-fMRI during a bilateral finger-tapping motor task (n = 11 subjects). Isotropic ADC-fMRI spatial maps show a group-level activation cluster overlapping with the left primary motor cortex and left primary somatosensory cortex (Figure 4A and Figure S14). Observing the overlaid subject-level spatial maps following subject-level cluster correction (Figure 4B) shows that bilateral activation was detected across individuals. Spatial maps from b200-dfMRI, b1000-dfMRI and BOLD-fMRI showed group-level widespread bilateral activation of the motor cortices, and BOLD-fMRI also detected activation in the basal ganglia (Figure 4C and Figure S12). There was also some activation in the temporal lobe, which may reflect activation of brain areas involved in interpreting the task instructions [40]. For completeness, BOLD-fMRI group-level spatial maps are shown when convolving the task timeseries with the haemodynamic response function (Figure S12), and ADC-fMRI group-level spatial maps are shown when convolving the task timeseries with the diffusion response function [17] (Figure S13).

**Figure 4:**
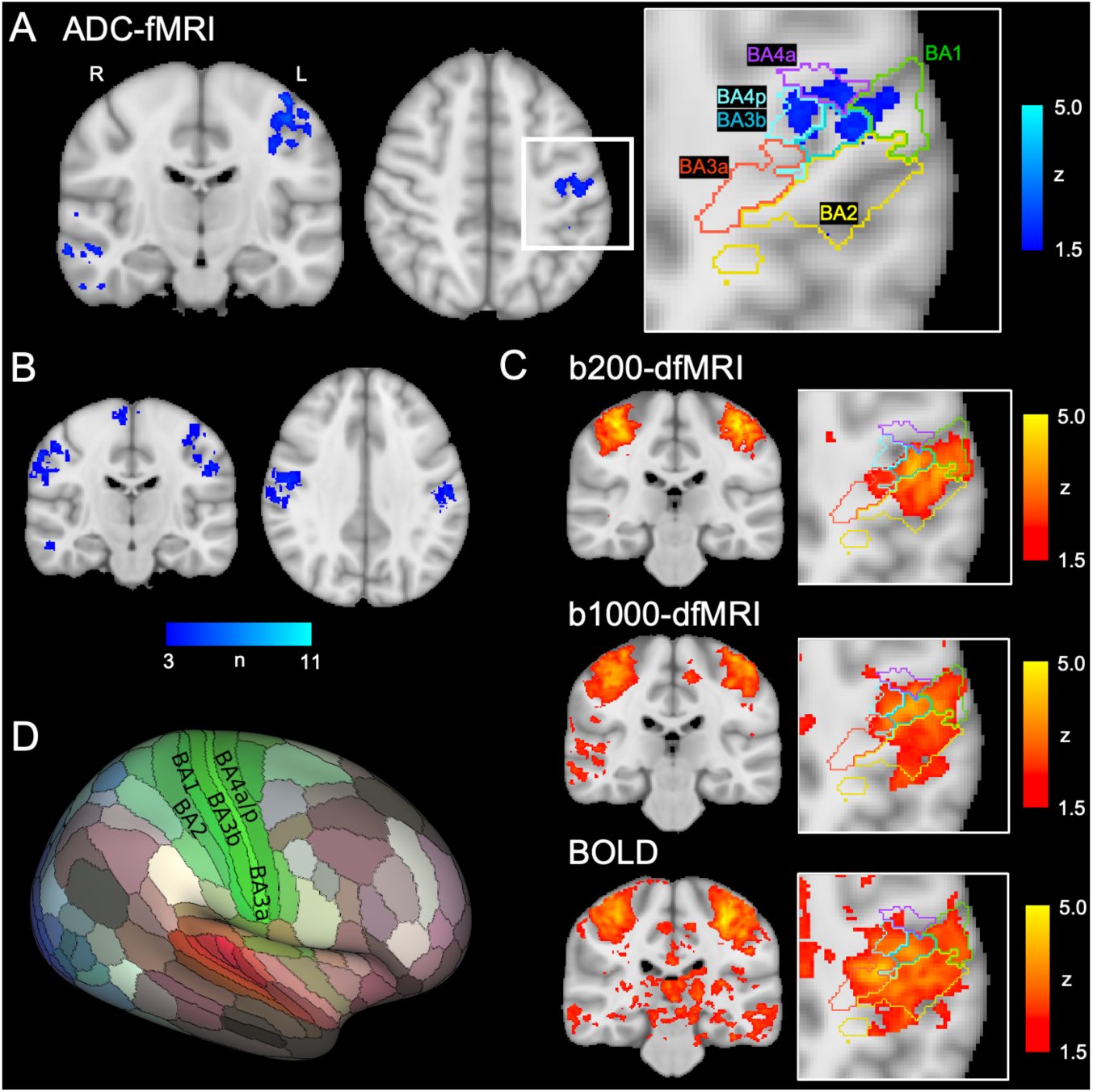
Motor task spatial maps. Spatial maps are shown for isotropic ADC-fMRI following group-level cluster correction (*z* 2’: 1.5, p *<* 0.05) with a zoomed-in panel showing Brodmann Areas (BA) defining the primary somatosensory cortex (BA1, 2, 3a, 3b) and primary motor cortex (BA4a, 4p), delineated by the Juelich atlas (A). Subject-level activation maps (following cluster correction at *z* 2’: 2.3, p *<* 0.05; shown in Figure S15) are aggregated for isotropic ADC-fMRI (B), with the colourbar indicating the number of subjects (among n = 11) for which each voxel was detected as active. Spatial maps following group-level cluster correction (*z* 2’: 1.5, p *<* 0.05) are also shown for b200-dfMRI, b1000-dfMRI and BOLD-fMRI (C). For anatomical reference, BA1-4 are displayed on an inflated brain surface (D) [44].

Spatial maps from b200-dfMRI, b1000-dfMRI and BOLD-fMRI were spread across Brodmann Areas (BA) delineating primary somatosensory cortex (BA1, BA2, BA3a, BA3b) and the posterior part of the the primary motor cortex (BA4p). ADC-fMRI activation was more spatially specific to the anterior and posterior parts of the primary motor cortex (BA4a, BA4p), and surrounding somatosensory regions BA1 and BA3b, around the area associated with hand movements [41–43].

Similar to the visual response, ADC-fMRI detected a larger proportion of active voxels in white matter than dfMRI and BOLD-fMRI. In subject-level cluster-corrected spatial maps from the motor task, isotropic ADC-fMRI detected 41.7% of voxels in white matter compared to 24.0% with b200-dfMRI, 23.1% with b1000-dfMRI, and 23.1% with BOLD-fMRI. Comparison of these proportions at the subject level confirmed significant differences between contrasts (p*<* 0.05; Figure S7, Table S7).

The temporal characteristics of the motor response (Figure 5) reflect those observed in the visual response, with ADC-fMRI exhibiting sharp onset and return to baseline (Table S2). BOLD-fMRI also had similar rise and fall times (4.4 and 6.0 s) to those in the visual task. However, b200-dfMRI and b1000-dfMRI had a quicker decay from task offset to 50% activation in the motor task (2.6 and 1.3 s respectively) than in the visual task. This appears to be due to an initial steep drop upon ending the task, followed by a longer tail (below 50% activation) resembling the BOLD delay. Comparison at the subject level confirmed that these differences between contrasts were significant (p *<* 0.05; Figure S8, Table S4). The peak amplitude (Table S1) of the BOLD-fMRI response to the motor task was much smaller than for visual stimulation (1.1% vs 2.5%), whereas the ADC-fMRI response was similar in both tasks (-1.3% vs -1.1%).

**Figure 5:**
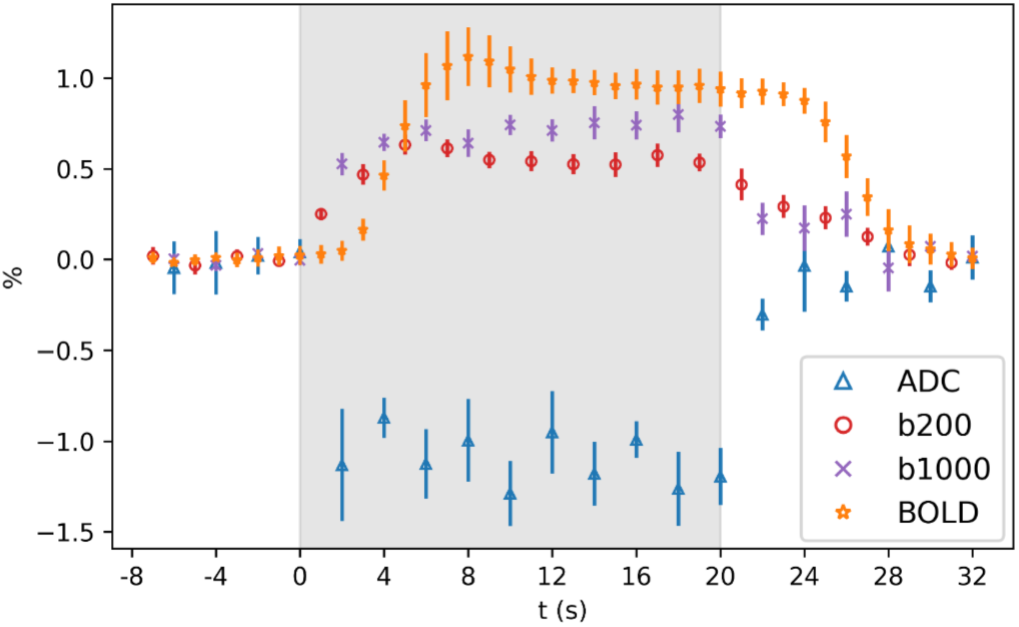
Motor task response in active voxels from subject-level spatial maps. Plots show the average timecourse, with bars indicating the standard error across subjects (n = 11). Subject-specific timecourses were averaged across epochs and across voxels in spatial maps following subject-level cluster correction (*z* 2’: 2.3, p *<* 0.05). Time is given in reference to the task onset, with the task stimulation duration indicated by the shaded area. Responses in grey and white matter voxels are shown in Figure S16.

## Discussion

In this study, we introduced a spherical b-tensor encoding ADC-fMRI acquisition for detection of neural activity in grey and white matter, independent of underlying tissue organisation. To achieve the same directional independence with linear diffusion encoding would require acquisition of at least three orthogonal directions, thus we provide a three-fold improvement in temporal resolution. To suppress vascular contributions to the signal, we used a shifted ADC approach calculated with b-values of 200 and 1000 s mm^-2^ [23, 25, 27, 28]. We demonstrated that isotropic ADC-fMRI detects neural activity at the group level during separate visual and motor tasks, with higher temporal specificity than dfMRI and BOLD-fMRI. In white matter, isotropic ADC-fMRI overcame the sensitivity to fibre direction exhibited by linear ADC-fMRI and the limited sensitivity exhibited by BOLD-fMRI. Thus, isotropic ADC-fMRI provides a method for detection of neural activity which is unbiased in terms of: i) brain tissue type; and ii) axonal/dendritic organisation at the voxel-level.

There is continued controversy surrounding the white matter BOLD signal, with many earlier studies simply disregarding it as a nuisance regressor due to a lack of clear evidence supporting neural origins of the signal [4]. More recent work suggests that the BOLD signal in white matter does reflect neural activity, however the reduced vasculature and altered haemodynamic response require distinct analysis methods to detect white matter activation [5–7], with task-dependent activation often only becoming apparent when averaging the BOLD signal within a region of interest. Additionally, the shape of the haemodynamic response function has been shown to vary with white matter depth [7], obstructing the use of voxel-wise general linear model analysis with BOLD-fMRI. This is apparent in our results from visual and motor tasks; BOLD-fMRI spatial maps showed primarily grey matter activation, despite using a multi-echo acquisition to estimate *T* ^⇤^ in each voxel [45], making the BOLD acquisition robust to the different *T*_2_ of grey and white matter [5]. The BOLD-fMRI response in white matter was primarily subcortical and had lower amplitude than in grey matter, possibly indicating partial volume effects with grey matter. Thus, even though it may be possible to detect white matter activity [6, 7, 37, 46, 47], BOLD-fMRI still expresses a strong preference for detection of grey matter activity. Results of our model-free frequency analysis of BOLD-fMRI data, following methods in [37], exemplify these limitations; while BOLD-fMRI is able to detect activity in white matter at the task frequency, this activity has a different response in different brain regions due to the heterogeneity of the haemodynamic response [7]. The white matter BOLD response is slower than that of grey matter (which is already delayed with respect to the underlying neuronal activity), which obscures simultaneous detection of grey and white matter activity, as demonstrated by the low sensitivity to white matter activation in voxel-wise general linear model analysis. This also obstructs analyses which assume covarying BOLD activity arises from simultaneous neural activity, such as resting-state functional connectivity [24].

The microscopic neuromorphological alterations which are assumed to cause measurable changes to ADC occur in myelinated axons [16] in addition to cell bodies [12, 13], neurites [13, 14], synaptic boutons [14] and astrocytic processes [15]. This may explain why ADC-fMRI expressed no clear preference for detecting activity in grey or white matter voxels. In line with previous findings of activity-related changes to diffusivity [48], we found comparable magnitude of ADC changes in visual and motor regions, as well as in both grey and white matter. Thus, while the heterogeneity of the BOLD response obstructs simultaneous mapping of activity in grey and white matter, ADC-fMRI provides brain-wide mapping of activity, lending it to whole-brain functional connectivity studies including white matter regions (e.g. resting-state ADC-fMRI [24]). In white matter, ADC changes in a voxel may be influenced by both the number and synchrony of action potentials. It may be possible that there is some coherence of action potentials along the length of the tract, theoretically allowing the direction of information propagation to be traced. However, this would occur on a timescale much smaller than that of our TR. Therefore, the measured morphological changes at the spatial and temporal scale in this study most likely represent an overall increase in the number of action potentials underlying block-design sustained visual or motor activity.

Due to the directional organisation of white matter fibres, activity-induced microstructural changes result in larger percentage change in water diffusivity perpendicular to fibres, compared to parallel, as demonstrated in vivo in the mouse optic nerve [29]. When measured with linear diffusion encoding gradients, the measured ADC change is therefore dependent on the angle between the white matter fibres and the diffusion encoding direction, as demonstrated by our in silico experiments. Previous studies have overcome this directional dependence by averaging over multiple volumes with different diffusion encoding directions [21, 22, 31], however this greatly reduces the temporal resolution. Spherical b-tensor encoding provides a method of measuring average ADC over all directions per signal acquisition. This has previously been used in a preclinical dfMRI study of somatosensory activation in mice [49]. Their analysis focused on the comparison of grey matter activation between isotropic dfMRI and BOLD-fMRI, and as such did not assess directional dependence in white matter, or compare with linear encoding. Furthermore, Nunes et al. [49] analysed isotropic diffusion-weighted timecourses rather than ADC timecourses, retaining T_2_-weighting and thus BOLD contributions. In our study, we used isotropic ADC-fMRI on a clinical scanner, showing that we overcome the dependence on directionality observed with linear encoding, and the dependence on BOLD contributions. By measuring the underlying fibre orientation in white matter, we demonstrated that the magnitude of ADC change was independent of the angle between the fibre direction and the reference direction for b-tensor encoding. Therefore, in addition to providing unbiased mapping of activity across grey and white matter, ADC-fMRI also provides unbiased mapping of activity across all white matter areas, independent of fibre organisation. We note for completeness that the spherical diffusion encoding effectively weights the signal by the trace of the diffusion tensor in the case when diffusion displays no time-dependence. In case of diffusion time-dependence, the estimated ADC may exhibit both an overall bias (which is not problematic for fMRI where we examine relative changes between baseline and task) and some orientation-dependence [50]. However, this effect was estimated to be relatively small (up to 5%) [50] and further minimal in the case of clinical diffusion times (50–80 ms), as diffusion time-dependence has been demonstrated in brain tissue in vivo either for short times (*<*40 ms) [51–53] or very long times (*>*100 ms) [54]. Based on our waveform we estimated the diffusion times of this sequence to be from 30 ms up to a limit imposed by the encoding duration (*<*80 ms).

The primary motivation for developing an alternative functional contrast to BOLD is to overcome the reliance on neurovascular coupling. Many prior studies have used diffusion-weighted spin-echo timecourses to remove signal contributions from intravascular frequency shift and extravascular static dephasing [55], detecting changes in diffusivity with earlier onset than BOLD-fMRI [11, 17–20, 56, 57]. Preclinical results have shown, using neurovascular coupling inhibition, that the dfMRI signal is dependent on neural activity via mechanisms other than neurovascular coupling [58, 59]. However, hyperoxia and hypercapnia experiments, which aim to trigger a vascular response in the absence of neural activity, showed that the BOLD response was attenuated in diffusion-weighted timecourses but not completely eliminated [20, 58]. The dfMRI signal remains sensitive to *T*_2_ changes in the extravascular tissues associated with deoxy-haemoglobin concentration (mainly around capillaries, therefore not refocused by spin-echo sequences), causing delayed return to baseline [17, 20]. This is evident in our results from b200-dfMRI and b1000- dfMRI which exhibit earlier onset than BOLD-fMRI (via diffusion-weighting) but slow return to baseline (via *T*_2_-weighting) (Figures 2 and 5). Using ADC-fMRI, these contributions to the signal are eliminated by taking the ratio between signals at different b-values to calculate ADC [19, 21, 22, 31, 49]. In contrast with many previous ADC-fMRI studies [19, 21, 49], we also minimise direct contribution to the contrast from blood water by adopting a shifted ADC approach [27, 28], calculating the ADC from b-values *>*200 s mm^-2^ [22, 23, 25].

Interactions between diffusion-weighting gradients and the susceptibility-induced background field gradients, which vary around vessels during the haemodynamic response, introduce another potential link between the measured ADC signal and the vascular response [60]. In our linear ADC-fMRI acquisition, the twice-refocused spin-echo sequence minimises this modulation of effective diffusion-weighting. However, the isotropic ADC-fMRI sequence is not compensated for cross-terms, meaning there may be some contamination of the isotropic ADC-fMRI signal with the vascular response. This may explain the slightly delayed return to baseline in the visual task response with isotropic ADC-fMRI in comparison with linear ADC-fMRI (Figure 2). In the response to the motor task, where the signal from the vasculature is much smaller (as shown by the lower amplitude BOLD response) this delayed return to baseline is no longer visible. It is possible that this delayed return to baseline is partially due to a delayed astrocytic response [15, 27, 61]. However, this delay is not seen in the linear ADC-fMRI response, as is much less apparent in the isotropic ADC-fMRI response to the motor task, where the BOLD response is much lower in magnitude, suggesting it is more likely due to the residual BOLD contamination of the isotropic ADC-fMRI sequence. In future, the waveform used in the isotropic encoding sequence can be designed to compensate for cross-terms in order to further eliminate vascular contamination [62].

In the ADC-fMRI acquisitions, it is difficult to separate the effects of reduced sensitivity due to lower contrast to noise ratio and increased spatial specificity. The group-level isotropic ADC-fMRI spatial map from the motor task shows unilateral activation of the left primary motor and somatosensory cortices (Figure 4A), corresponding to areas associated with hand movement [41–43] in the hemisphere contralateral to the participants’ dominant hand. This was consistent with the peak z-score of the BOLD-fMRI spatial map being in the left hemisphere. While isotropic ADC-fMRI detected bilateral activation at the individual level (Figure 4B), the increased specificity (illustrated by the spatial overlap with the hand movement areas) but also poorer sensitivity as compared to BOLD-fMRI resulted in low overlap between spatial maps of individuals [63]. Therefore, with our sample size, only unilateral activation survived group-level cluster correction for the motor task. Conversely, the widespread vascular response detected by b200dfMRI, b1000-dfMRI and BOLD-fMRI gives more spatial overlap between subjects but also covers motor areas that are not specific to the hand. Although we cannot determine whether the smaller spatial extent of activation with ADC-fMRI is primarily due to increased specificity or decreased sensitivity, Figure 1 shows that ADC-fMRI was able to detect areas of activation in the optic radiation in response to visual stimulation, which were not detected by BOLD-fMRI. Thus, even despite the reduced sensitivity of ADC-fMRI, we have shown that it can detect activation in the white matter that is otherwise not mapped by BOLD-fMRI.

The activity detected in the optic radiation appears to be isolated to the posterior regions of the tract, rather than distributed along its entire length. Similarly, activity detected in the corticospinal tract was limited to areas close to the cortex. This could be due to microstructural variations along the tract, such as a distribution in axon diameters, which may induce a difference in both the coherence of action potentials and the relative intra- and extra-cellular volume change resulting from a given deformation during action potential firing, ultimately inducing different magnitude ADC changes. Alternatively, this may simply be due to the lower SNR deeper in the brain obscuring detection of activity (in isotropic ADC-fMRI data for the visual task, temporal SNR was around 15 near the thalamus, compared to around 30 near the cortex). Theoretically, the use of higher b-values would increase sensitivity to diffusion changes [17], possibly allowing detection of activation along the full optic radiation. However, the even lower SNR associated with higher b-values makes them practically less favourable for ADC-fMRI (we measured a temporal SNR of *<*15 throughout the optic radiation in linear ADC-fMRI data with b-values of [1000, 2000] s mm^-2^ and did not find any activation clusters in these data - see Supplementary Materials “High b-value Acquisition and SNR Comparison”). Although isolated to the higher-SNR areas, ADC-fMRI was better able to detect activity in white matter tracts than BOLD-fMRI, which primarily detected grey matter regions. Thus, with future developments to hardware, acquisition or post-processing denoising to improve sensitivity deeper in the brain, isotropic ADC-fMRI may provide a method of comprehensive whole-brain mapping of grey and white matter activity.

Isotropic ADC-fMRI and linear ADC-fMRI both detected a higher proportion of white matter voxels than BOLD-fMRI, which predominantly detected grey matter activity. Interestingly, isotropic ADC-fMRI and linear ADC-fMRI detected similar proportions of grey and white matter voxels in response to the visual task, despite the directional independence of isotropic ADC-fMRI which would intuitively result in an increase in detection of white matter voxels. This could reflect a corresponding increase in the detection of grey matter voxels with isotropic ADC-fMRI due to some residual BOLD contamination of the signal (as described above), which would likely be stronger in grey matter due to the increased vasculature. An alternative explanation could be related to some orientation dependence of linear ADC-fMRI not only in white but also in grey matter due to alignment of cortical columns [64–66].

Across all angles, the spread of activation amplitudes was lower in magnitude with isotropic encoding than with linear encoding (Figure 3B). This trend can also be seen in the response (Figure 2), which shows a lower amplitude with isotropic encoding than with linear encoding. This reflects the fact that isotropic encoding weights the signal by the trace of the diffusion tensor, simultaneously capturing the large changes perpendicular to fibres and the relatively small changes parallel to fibres [29]. This reduction in sensitivity, in addition to the current SNR constraints, is a possible limitation of isotropic ADC-fMRI. In summary, isotropic ADC-fMRI provides a new method of mapping activity in the human brain. The dependence of BOLD-fMRI on haemodynamic coupling imposes limitations when mapping activity across white matter due to the heterogeneity of the haemodynamic response. By exploiting neuromorphological coupling, ADC-fMRI is able to detect activation in grey and white matter. Therefore, isotropic ADC-fMRI opens new avenues for unbiased investigation of brain-wide functional activation, such as monitoring functional recovery of white matter areas following damage (for example following a stroke), neurosurgical planning based on white matter functional activation maps combined with tractography, or resting-state functional connectivity within and between grey and white matter, which remains an ongoing challenge for functional neuroimaging [24, 47, 67].

## Methods

This study was approved by the ethics committee of the canton of Vaud, Switzerland (CER-VD). All participants provided written informed consent. In the visual task, 13 healthy adults were included (age 21–37 years, median 27; 7 female). In the motor task, 11 healthy adults were included (age 21–29 years, median 23; 4 female; 1 left-handed).

### MRI Acquisition

MRI data were acquired using a 3T Siemens Magnetom Prisma with 80 mT/m gradients and 200 T/m/s slew rate, and a 64-channel head coil.

Whole-brain T1-weighted anatomical images were acquired, for anatomical reference and parcellation, using 3D Magnetization Prepared 2 Rapid Acquisition Gradient Echoes (MP2RAGE) [68] with the following parameters: 1 mm^3^ isotropic voxels; 256 x 256 mm^2^ field of view; 176 slices; repetition time (TR) 5000 ms; echo time (TE) 2.98 ms; inversion times (TI) 700, 2500 ms; flip angles 4°, 5°; in-plane acceleration factor 3 using Generalized Autocalibrating Partially Parallel Acquisitions (GRAPPA) [69].

Multi-shell diffusion-weighted imaging (DWI) data were acquired for estimation of fibre orientation, using a 2D multi-slice spin-echo echo planar imaging (EPI) sequence with the following parameters: (2 mm)^3^ isotropic voxels; 232 x 232 mm^2^ field of view; 60 slices; TR 5000 ms; TE 80 ms; in-plane acceleration factor 2; multiband factor 2 [70, 71]; anterior-posterior phase encoding. Three b-values were acquired: b = 1000, 2000, and 3000 s mm^-2^ with 20, 30, and 48 directions respectively. Diffusion encoding directions on each b-value shell were equally distributed according to electrostatic repulsion. Four interspersed b = 0 volumes were acquired. Two additional b = 0 volumes were acquired with reverse EPI phase encoding direction (posterior-anterior) for correction of *B*_0_ field inhomogeneity distortions.

Isotropic dfMRI data were acquired with a spherical b-tensor encoding sequence [34], and linear dfMRI with a twice-refocused spin-echo EPI sequence with bipolar linear encoding gradients. The spherical btensor encoding waveform was numerically optimised and compensated for concomitant gradients. DfMRI acquisitions used the following parameters: (2.5 mm)^3^ isotropic voxels; 50% slice gap; 232 x 232 mm^2^ field of view; 16 slices; TR 1000 ms; TE 82 ms (isotropic dfMRI, visual task), 84 ms (isotropic dfMRI, motor task), 72 ms (linear dfMRI); flip angle 90°; in-plane acceleration factor 2; partial Fourier factor 6/8; multiband factor 2. During dfMRI acquisitions, two b = 0 volumes were acquired before the start of the task, followed by alternating volumes at b = 200 and 1000 s mm^-2^ for the duration of the task. The low b-value (200 s mm^-2^) was selected to maximise signal whilst minimising contributions from the blood water signal (see Introduction). The high b-value (1000 s mm^-2^) was selected to maintain reasonable SNR. For linear dfMRI, a single diffusion encoding direction was chosen for all volumes. This was defined as equal x, y, z weighting.

To enable a short TR = 1 s, the imaging volume gave partial brain coverage. For the visual task, this slab was aligned in axial plane to encompass the visual cortex and optic radiation. For the motor task, the slab was aligned in the coronal plane to encompass the motor cortex and corticospinal tract. Anterior-posterior phase encoding was used for the visual task and left-right phase encoding used for the motor task. For each acquisition, two additional (b = 0) volumes were acquired with reverse phase encoding for correction of *B*_0_ field inhomogeneity distortions.

Multi-echo BOLD-fMRI data [72] were acquired with a gradient echo sequence with four echoes (TE 12.60, 33.22, 53.84, 74.46 ms) with flip angle 62°. All resolution, brain coverage and acceleration parameters were matched to the dfMRI acquisitions. Two additional volumes were acquired with reverse phase encoding for correction of *B*_0_ field inhomogeneity distortions.

### Task Paradigm

Visual stimuli/cues for the visual and motor tasks were generated using PsychoPy [73]. The visual task consisted of a block design, alternating between stimulus periods displaying an 8 Hz flashing radial checkerboard, and rest periods displaying a fixation cross. Stimulus blocks were 12 s, and were separated by 18 s rest blocks. One full visual stimulus paradigm consisted of 16 stimulus epochs with four interspersed 30 s baseline blocks, for a scan duration of 10 minutes. The motor task consisted of 20 s self-paced fingertapping blocks, cued by instructions on the screen, separated by 20 s rest blocks displaying a fixation cross, repeated 15 times for a scan duration of 10 minutes. During finger-tapping blocks, participants were instructed to sequentially tap each finger against their thumb, from the first finger to the little finger and back again, with both hands.

### Preprocessing

#### DfMRI

Magnitude image denoising was applied separately to the b = 200 s mm^-2^ timeseries and the b = 1000 s mm^-2^ timeseries using NORDIC [74], with a kernel size 7x7x7 and step size 1 for both g-factor estimation and PCA denoising. Following Gibbs unringing of all volumes with MRtrix3 [75, 76], FSL Topup was used to estimate the susceptibility bias field from b = 0 volumes, which was then used to correct susceptibility-induced distortions in all volumes [77, 78]. ANTs motion correction was applied to each b-value timeseries separately, which were then registered to the initial b = 0 volume [79]. A mask of brain tissue was created by applying Synthstrip to the Topup-corrected b = 0 volumes [80] and used to remove non-brain voxels from the full timeseries.

#### BOLD-fMRI

ANTs motion correction was calculated on the timeseries of the first echo of the multi-echo BOLD data, then these transformations were used to correct the timeseries for all echoes. Tedana was used to calculate an optimally combined signal from a weighted average of echoes (including denoising with TEDPCA and TEDICA) [45]. The susceptibility bias field was calculated from the first echo using FSL Topup, then used to correct distortions in the optimally combined data. A brain mask was created using Synthstrip to remove non-brain voxels.

#### DWI

MP-PCA denoising was applied to DWI data [81–83], with patch size 5x5x5, followed by Gibbs unringing and FSL Topup. Normalised fibre orientation distribution (FOD) images were obtained using MRtrix3 by deconvolving the estimated response function [84] from the data using multi-tissue constrained spherical deconvolution [85].

#### T1 Images

T1 images were denoised using spatially adaptive non-local means filtering with ANTs [86], then skullstripped with Synthstrip [80] and segmented into tissue maps using FSL Fast [87]. Grey and white matter masks were transformed to the image space of each functional acquisition (dfMRI and BOLD-fMRI) using rigid-body registration with ANTs in order to assess the activation in grey and white matter. Freesurfer was used to create subject-specific masks of the ventricles [88] to allow the ventricle signal to be included as a covariate in task analysis (see below). For group-level analysis, T1 images were nonlinearly registered to MNI standard space using ANTs.

### Task Activation

We investigated task-based activation in BOLD-fMRI, and both linear and isotropic ADC-fMRI, b200- dfMRI and b1000-dfMRI timeseries using FSL FEAT (FMRI expert analysis tool) version 6.00 [89]. Following highpass temporal filtering to correct for signal drift (100 s cutoff), a general linear model was used to measure association between the timeseries and the task, with the average ventricle signal included as a covariate. To avoid any assumptions about the shape of the response function, we modelled the task as a boxcar function for all contrasts. A dfMRI response function had been previously derived which aims to distinguish the diffusion response from the haemodynamic response [17], however this was calculated from dfMRI data, not ADC-fMRI, therefore some haemodynamic contribution to the diffusion response cannot be entirely ruled out. For completeness, supplementary results are presented for ADC-fMRI with the task modelled as a boxcar function convolved with the diffusion response function [17]. Additionally, supplementary results are presented for BOLD-fMRI with the task modelled as a boxcar function convolved with the canonical haemodynamic response function. We investigated negative association with ADC-fMRI and positive association with all other timeseries. Subject-level spatial maps were cluster-corrected at *z* 2’: 2.3, p *<* 0.05. Group-level analysis was carried out using FLAME (FMRIB’s Local Analysis of Mixed Effects) stage 1 and 2 [90], with cluster correction at *z* 2’: 1.5, p *<* 0.05.

### Response

To plot the task response of each contrast, the functional timeseries was averaged across voxels in the subject-level cluster-corrected spatial map. Each epoch was then normalised to its baseline, and this was averaged across epochs. To characterise the response timings of each contrast, the response was averaged across subjects and linearly interpolated to a timestep of 0.1 s. For b200-dfMRI acquisitions, where the acquisition timings did not fall exactly on the start and end times of the task, it was assumed that the signal at these time points was equal to the previously acquired volume. The time to reach 50% of peak activation amplitude was then measured from the interpolated timeseries.

### Sensitivity to Fibre Direction

Fibre direction was defined as the direction of the largest FOD peak, and was measured in each white matter voxel from the FOD image for each subject. This was transformed to the image space of each functional acquisition in order to measure the angle between the fibre direction and the diffusion encoding direction for linear ADC-fMRI, or spherical b-tensor encoding reference direction for isotropic ADC-fMRI. This angle was measured in each active voxel in subject-level cluster-corrected spatial maps, and the distribution of fibre angles was compared between linear and isotropic ADC-fMRI using a two-sample Kolmogorov-Smirnov test. To plot the fibre angle against activation magnitude, the activation of each voxel was measured as the percentage change in ADC during task compared to baseline, averaged over epochs. If the activation of a voxel was more than three standard deviations higher than the mean activation across all subjects, this voxel was deemed an outlier and rejected from this analysis.

### *In Silico* Experiments

Numerical phantoms were generated using the CATERPillar tool [39], designed to simulate white matter substrates. These phantoms featured parallel axons, with diameters drawn from a Gamma distribution with mean of 1 *µ*m, and exhibiting both beading and tortuosity, in a voxel size of (100 *µ*m)^3^. The axonal beading occurred periodically, with an amplitude equal to 0.3 times the average axon radius. The tortuosity, defined as the ratio of the total axon length to the straight-line distance between its starting and ending points, had an average value of 1.2 (Figure S9). A baseline phantom was generated with an intracellular volume fraction of 50%. To simulate the conditions of axons during neural activity, axons were subjected to degrees of swelling at 0.25%, 0.5%, 0.75%, and 1% of their original volume.

We measured ADC at each degree of swelling, as a percentage change from baseline, using the Monte Carlo Diffusion Simulator [38]. Diffusion simulations comprised 1 million random walkers diffusing over 40,000 steps, each with a duration of fl*t* = 2 ⇥ 10^-3^ ms and a step length fl*s* = 0.15 *µ*m, based on a free diffusion coefficient *D*_0_ = 2 *µ*m^2^ ms^-1^. To generate synthetic signals analogous to those obtained from linear ADC-fMRI, we used a pulse gradient sequence with a time between pulses (fl) of 50 ms and a gradient pulse width (*<*) of 16.5 ms. Additionally, we simulated isotropic ADC-fMRI using the same spherical b-tensor encoding waveform as used in the in vivo acquisition. For both linear and isotropic simulations, ADC was calculated from b-values of 200 and 1000 s mm^-2^, and *T_E_* was set to 67 ms. *T*_2_ decay was not accounted for in the simulation. Diffusion signals were computed from the accumulated diffusion phase. To investigate the effect of fibre angle on measured ADC changes, linear ADC-fMRI simulations were repeated with linear encoding in 21 different directions, isotropically distributed over a sphere. Isotropic ADC-fMRI simulations were computed using each of these directions as the reference direction for b-tensor encoding. For each simulation, the intracellular and extracellular signals were recorded separately in addition to the combined signal.

## Acknowledgements

We acknowledge the CIBM Center for Biomedical Imaging for providing expertise and resources to conduct this study. In particular, we would like to thank Jean-Baptiste Ledoux and Eleonora Fornari for assistance with MRI scanning. We would also like to thank Antoine Delattre-Klauser, Tobias Kober, Tom Hilbert, Gian Franco Piredda and Xavier Sieber for assistance with MRI sequences. This work was supported by ERC Starting Grant ‘FIREPATH’, SERI no. MB22.00032. IJ is supported by an SNSF Eccellenza fellowship no. 194260.

## Author Contributions

IJ conceived the ideas and supervised the project. AS and IJ designed the methodology. AS, JND and IdR acquired data. AS analysed the data. JND carried out in silico experiments. IdR and FS contributed to MRI sequences. AS drafted the original manuscript. All authors reviewed and edited the manuscript.

## Competing Interests

F.S. is co-inventor in technology related to this research and has financial interests in Random Walk Imaging AB. All other authors have no competing interests to declare.

## Data Availability Statement

Data will be made publicly available on publication.

## Notes

### Summary of Updates

Supplementary analysis added, including model-free frequency analysis and diffusion response function analysis.

