## Supplementary Materials for "Mapping grey and white matter activity in the human brain with isotropic ADC-fMRI"

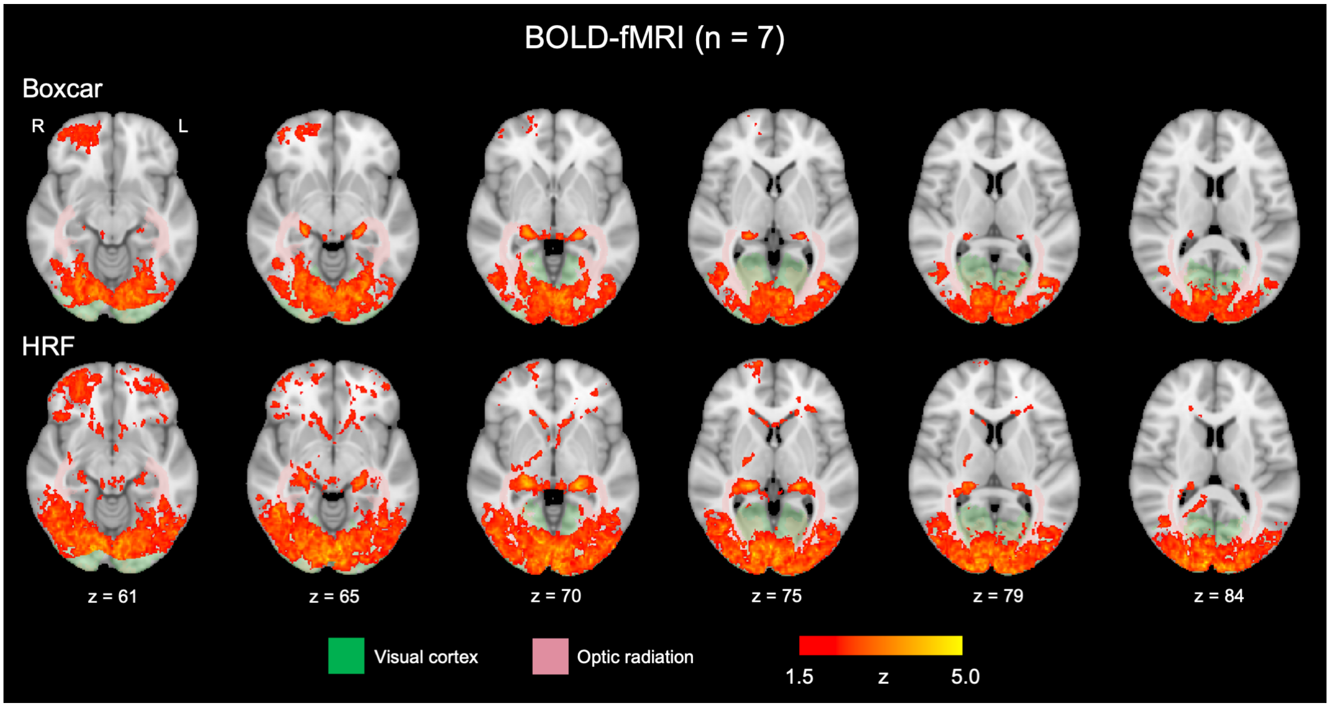


Figure S1: BOLD-fMRI spatial maps showing response to the visual task. The colour bar shows group-level z-score following group-level cluster correction ($z \geq 1.5$, $p < 0.05$). Spatial maps are shown for the response to the task modelled as a boxcar function (top), and modelled as the convolution of the boxcar function with the canonical haemodynamic response function (HRF; bottom). For anatomical reference, Juelich atlas regions defining the visual cortex (green) and optic radiation (pink) are overlaid on the MNI152 standard template.

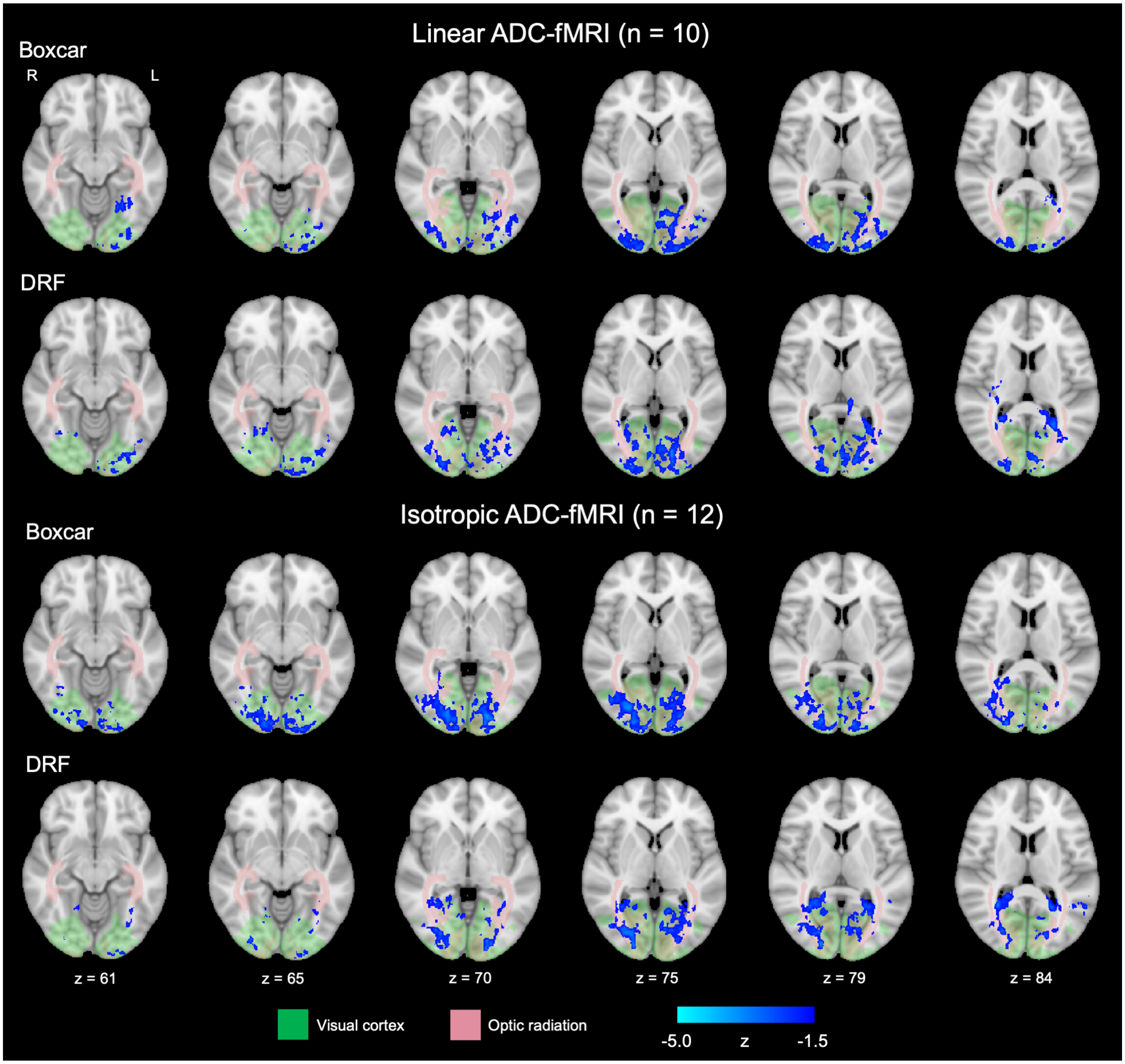


Figure S2: Linear and isotropic ADC-fMRI spatial maps showing response to the visual task with different response functions. The colour bar shows group-level z-score following group-level cluster correction ($z \geq 1.5$, $p < 0.05$). Spatial maps are shown for the response to the task modelled as a boxcar function, and modelled as the convolution of the boxcar function with the diffusion response function [17] (DRF). For anatomical reference, Juelich atlas regions defining the visual cortex (green) and optic radiation (pink) are overlaid on the MNI152 standard template.

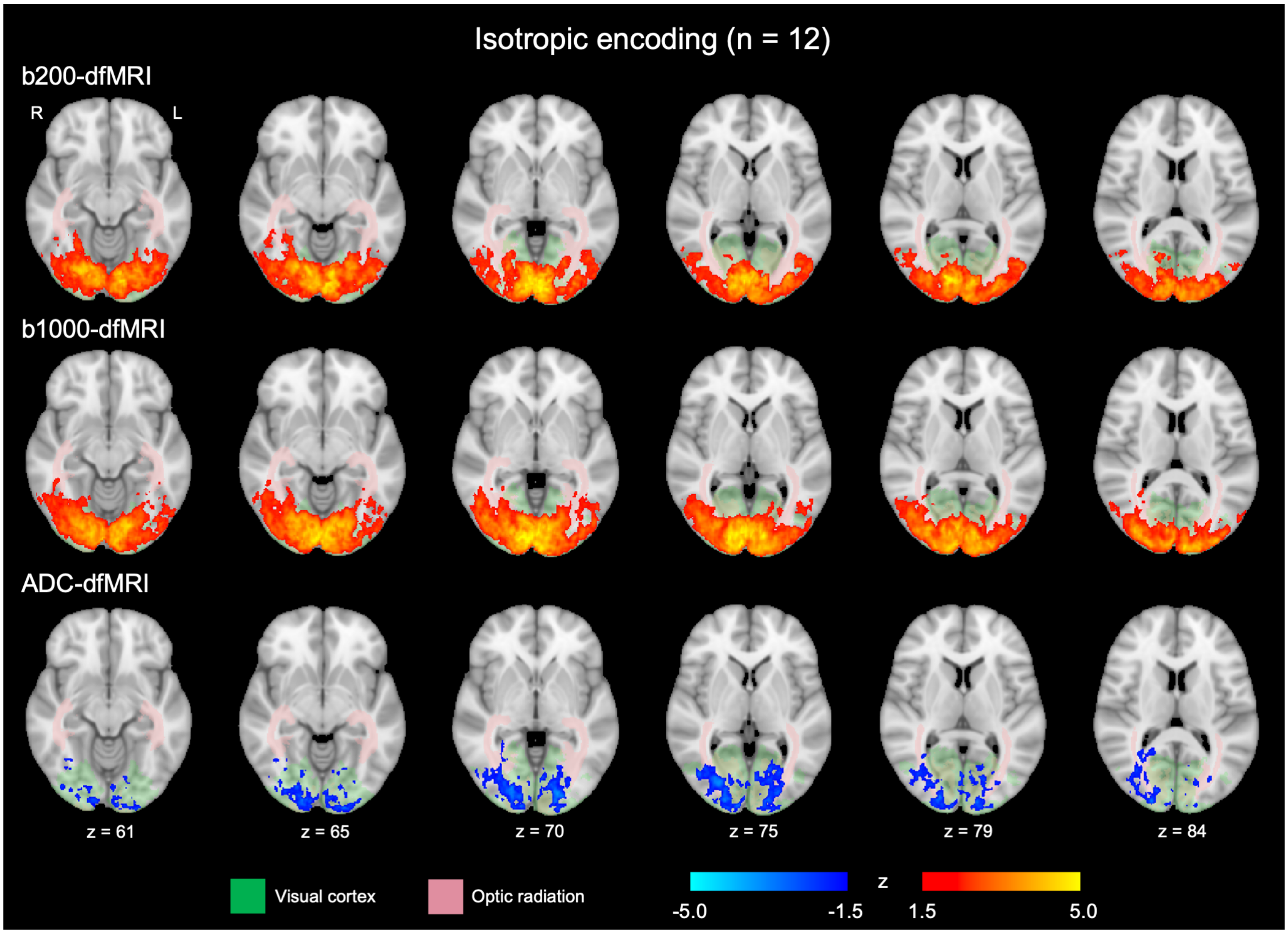


Figure S3: ADC-fMRI, b200-dfMRI and b1000-dfMRI spatial maps for isotropic encoding, showing response to the visual task. The colour bars show z-scores following group-level cluster correction ($z \geq 1.5$, $p < 0.05$). For anatomical reference, Juelich atlas regions defining the visual cortex (green) and optic radiation (pink) are overlaid on the MNI152 standard template.

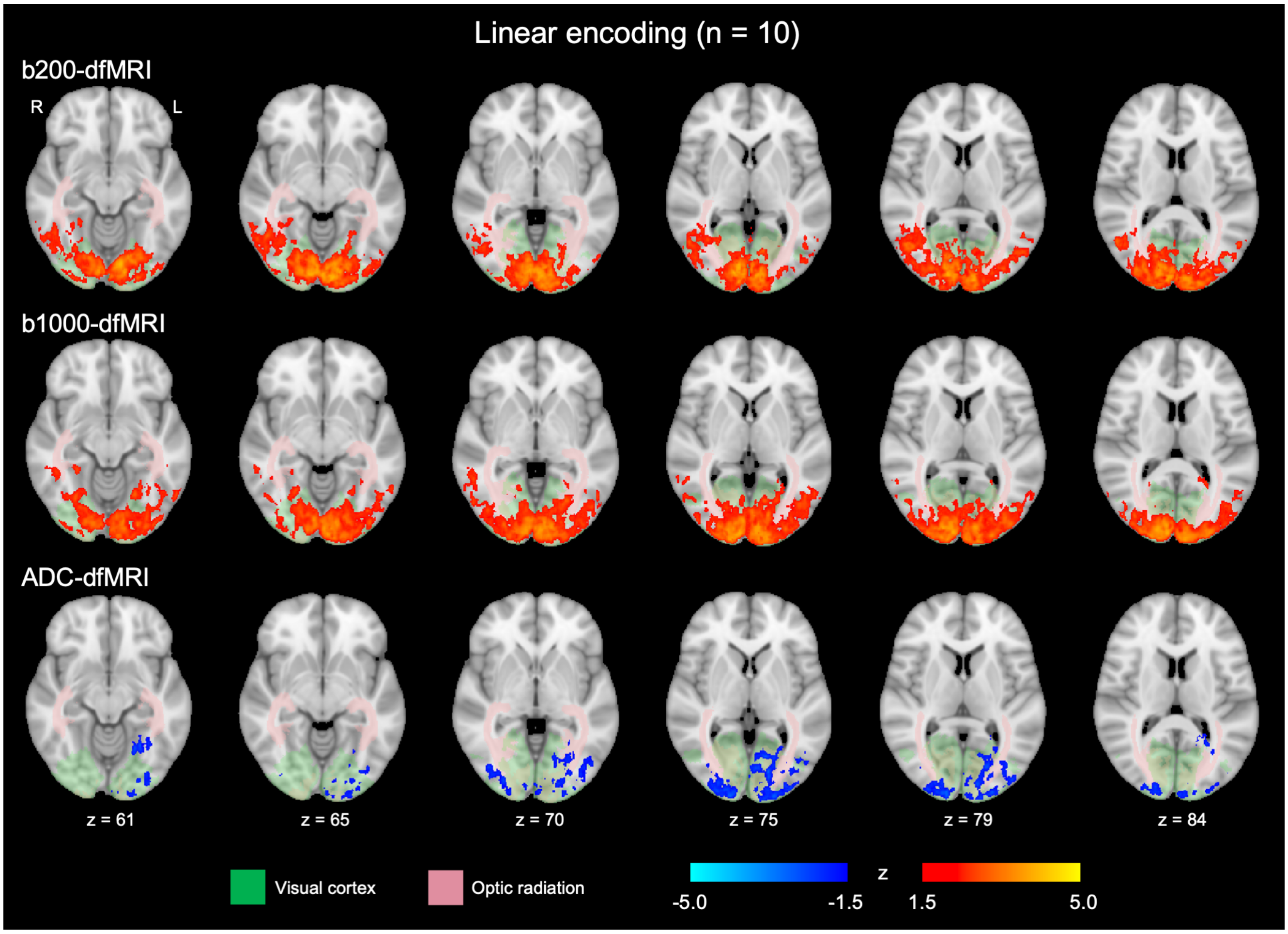


Figure S4: ADC-fMRI, b200-dfMRI and b1000-dfMRI spatial maps for linear encoding, showing response to the visual task. The colour bars show z-scores following group-level cluster correction ($z \geq 1.5$, $p < 0.05$). For anatomical reference, Juelich atlas regions defining the visual cortex (green) and optic radiation (pink) are overlaid on the MNI152 standard template.

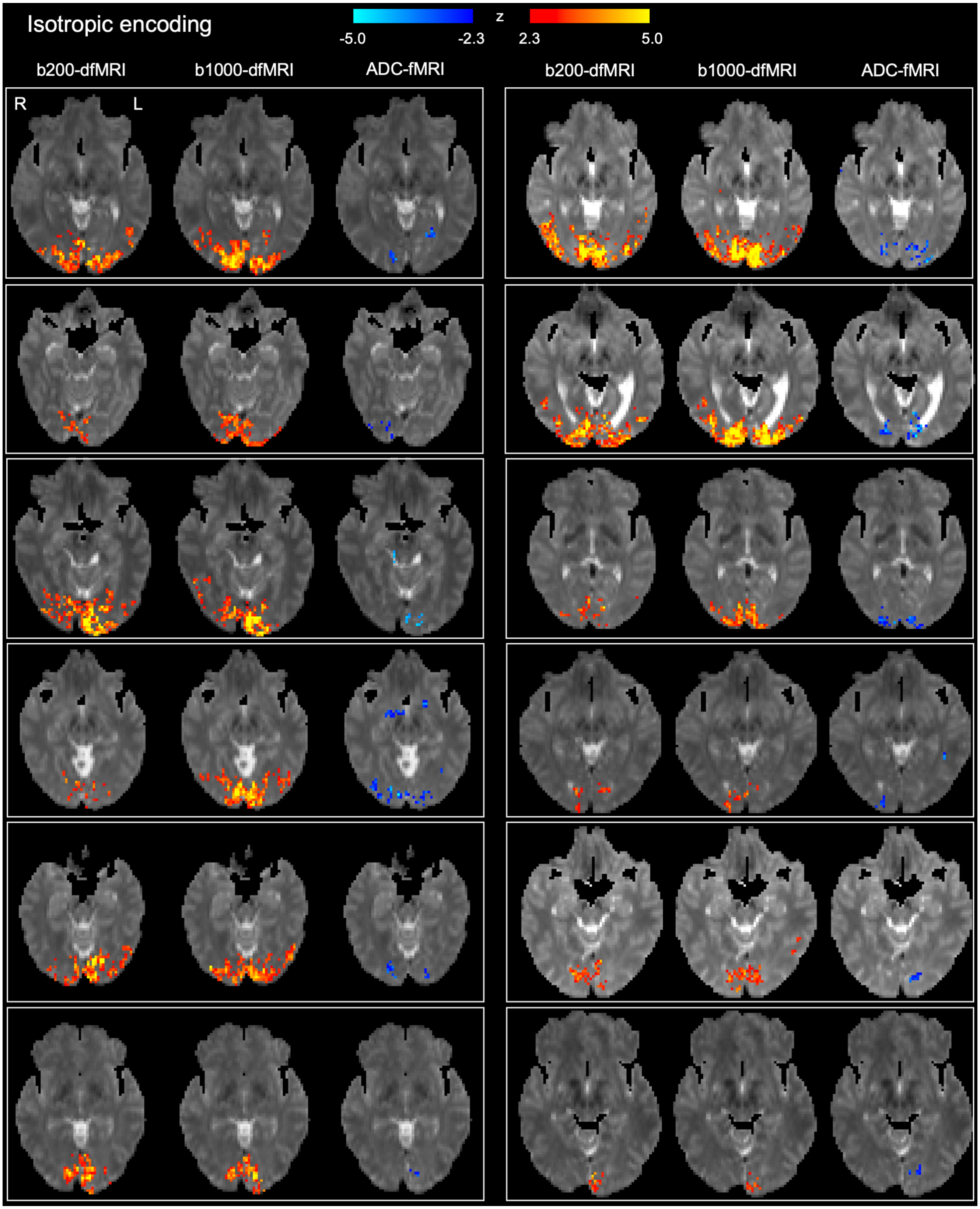


Figure S5: Subject-level visual stimulation spatial maps for isotropic encoding (n = 12). The colour bars show z-scores for b200-dfMRI, b1000-dfMRI and ADC-fMRI following subject-level cluster correction ($z \geq 2.3$, $p < 0.05$).

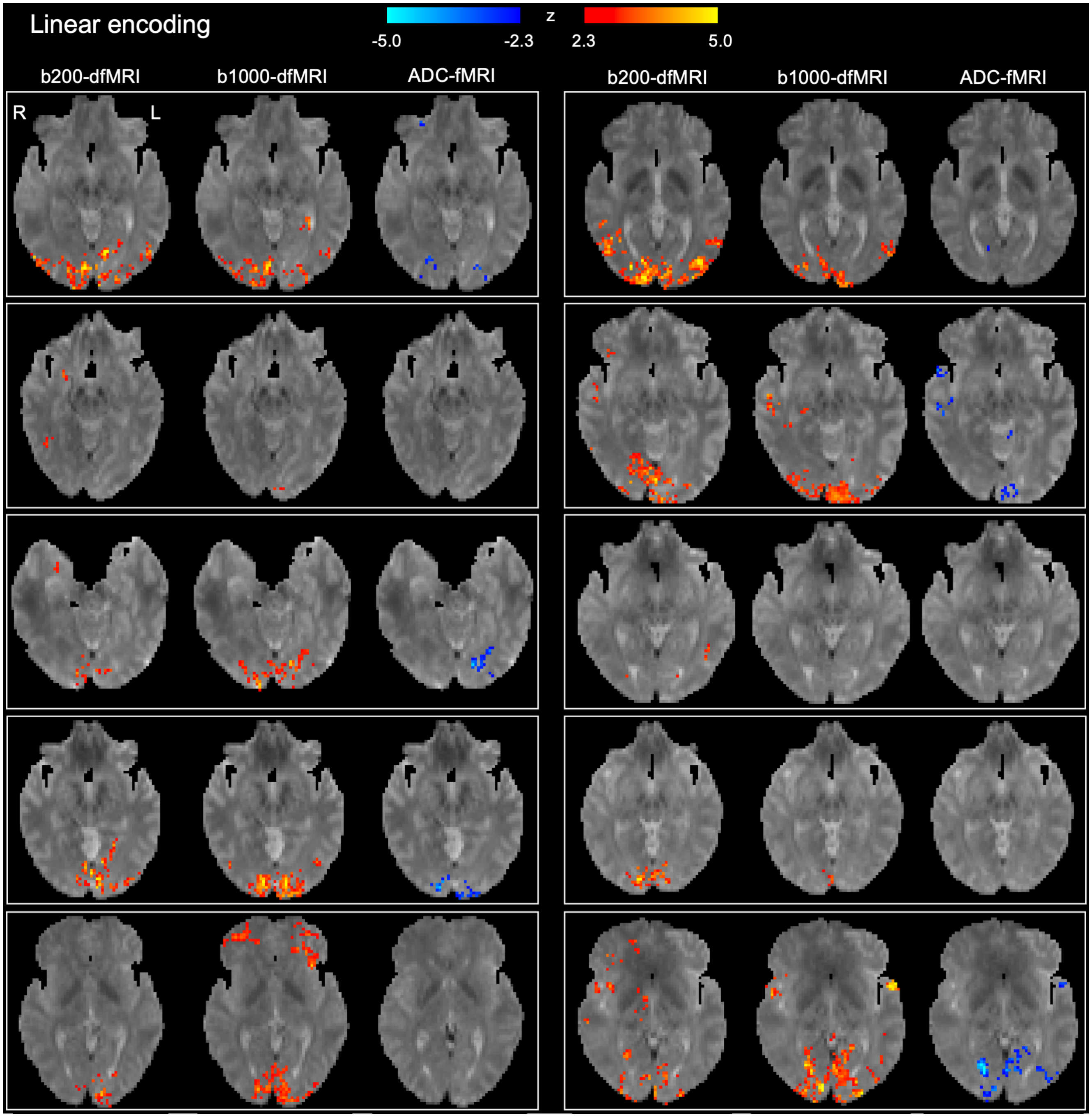


Figure S6: Subject-level visual stimulation spatial maps for linear encoding (n = 10). The colour bars show z-scores for b200-dfMRI, b1000-dfMRI and ADC-fMRI following subject-level cluster correction ($z \geq 2.3$, $p < 0.05$).

| Encoding | Contrast | All voxels (%) | Grey matter (%) | White matter (%) |
|---|---|---|---|---|
| **Visual Task** | | | | |
| Linear | ADC-fMRI | -1.3 | -1.3 | -1.3 |
| | b200-dfMRI | 0.7 | 0.7 | 0.5 |
| | b1000-dfMRI | 0.8 | 0.8 | 0.9 |
| Isotropic | ADC-fMRI | -1.1 | -1.2 | -1.0 |
| | b200-dfMRI | 1.0 | 1.1 | 0.7 |
| | b1000-dfMRI | 0.9 | 0.9 | 0.7 |
| | BOLD-fMRI | 2.5 | 2.7 | 1.2 |
| **Motor Task** | | | | |
| Isotropic | ADC-fMRI | -1.3 | -1.4 | -1.2 |
| | b200-dfMRI | 0.6 | 0.7 | 0.5 |
| | b1000-dfMRI | 0.8 | 0.8 | 0.7 |
| | BOLD-fMRI | 1.1 | 1.2 | 0.8 |

Table S1: Peak response amplitudes. For each contrast and acquisition, the time course was averaged across significant voxels in subject-level spatial maps (following cluster correction at $z \geq 2.3$, $p < 0.05$), then each epoch normalised to its baseline. Responses were averaged across epochs and subjects then the peak value of this average response was measured as a percentage change from baseline. This was repeated for significant voxels in grey and white matter.

| Encoding | Contrast | Rise time (s) | Fall time (s) |
|---|---|---|---|
| **Visual Task** | | | |
| Linear | ADC-fMRI | 1.3 | 1.1 |
| | b200-dfMRI | 1.4 | 4.3 |
| | b1000-dfMRI | 1.6 | 4.3 |
| Isotropic | ADC-fMRI | 1.5 | 1.8 |
| | b200-dfMRI | 2.6 | 4.7 |
| | b1000-dfMRI | 2.1 | 4.2 |
| | BOLD-fMRI | 4.1 | 5.7 |
| **Motor Task** | | | |
| Isotropic | ADC-fMRI | 1.2 | 1.2 |
| | b200-dfMRI | 1.7 | 2.6 |
| | b1000-dfMRI | 1.6 | 1.3 |
| | BOLD-fMRI | 4.4 | 6.0 |

Table S2: Response time characteristics. Time courses for significant voxels in subject-level spatial maps (following cluster correction at $z \geq 2.3$, $p < 0.05$) were averaged across epochs and subjects then interpolated to a time step of 0.1 s. Rise time and fall time of this average response were measured as the time from task onset or offset to 50% of the peak response amplitude (Table S1).

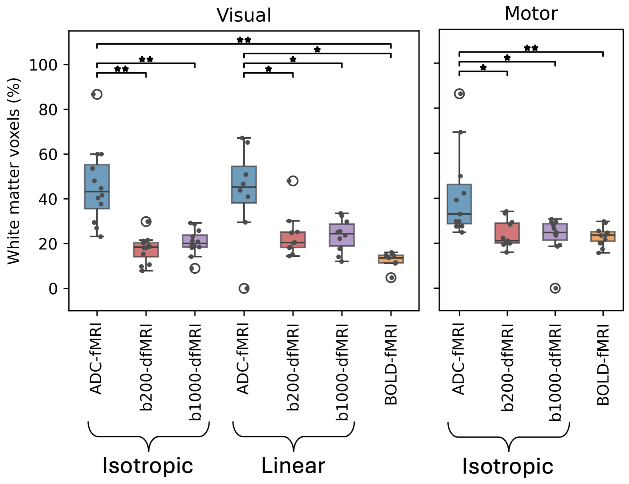


Figure S7: Comparison of the proportion of white matter voxels in subject-level cluster-corrected spatial maps. This was compared between contrasts using one-tailed Mann-Whitney U-tests with Bonferroni correction (to reduce multiple comparisons, comparisons between b200-dfMRI, b1000-dfMRI and BOLD-fMRI were not tested). *$p < 0.05$, **$p < 0.001$. Cohen's d effect sizes and p-values are shown in Table S3.

| | Isotropic | | Linear | | |
|---|---|---|---|---|---|
| | b200-dfMRI | b1000-dfMRI | b200-dfMRI | b1000-dfMRI | BOLD-fMRI |
| **Visual Task** | | | | | |
| Isotropic ADC-fMRI | 2.27 (0.0001) | 2.03 (0.0002) | | | 2.77 (0.0007) |
| Linear ADC-fMRI | | | 1.25 (0.0400) | 1.30 (0.0233) | 2.14 (0.0210) |
| **Motor Task** | | | | | |
| Isotropic ADC-fMRI | 1.27 (0.0087) | 1.28 (0.0047) | | | 1.37 (0.0010) |

Table S3: Cohen's d effect sizes and Bonferroni-corrected p-values, displayed as d (p), for the comparison of the proportion of white matter voxels in subject-level cluster-corrected spatial maps. Comparisons were made between contrasts using one-tailed Mann-Whitney U-tests. To reduce multiple comparisons, comparisons between b200-dfMRI, b1000-dfMRI and BOLD-fMRI were not tested.

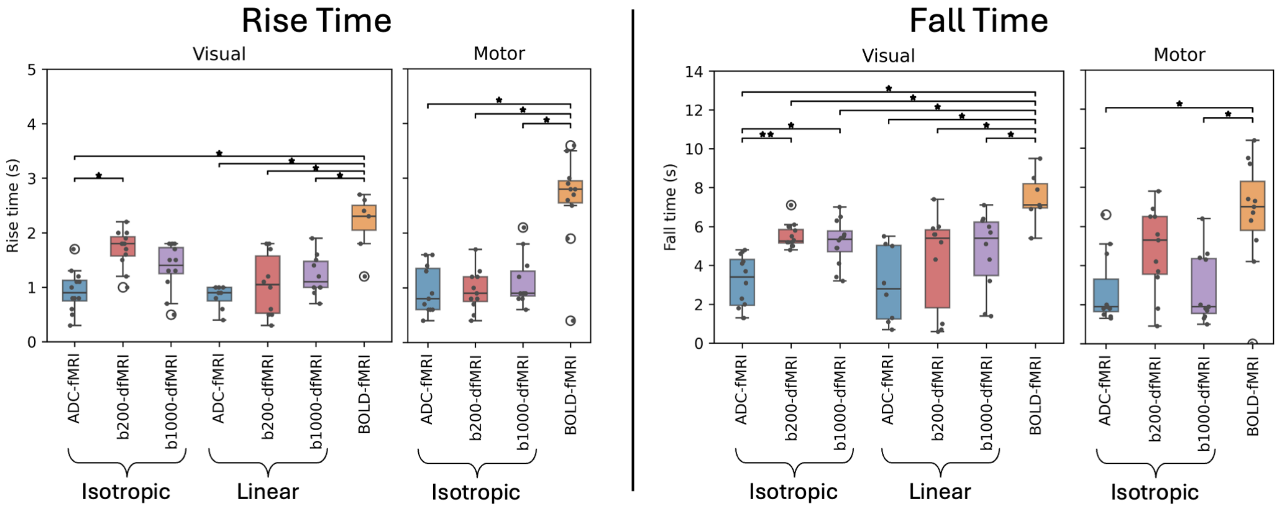


Figure S8: Comparison of subject-level response time characteristics across contrasts. Time courses for significant voxels in subject-level spatial maps (following cluster correction at $z \geq 2.3$, $p < 0.05$) were averaged across epochs then interpolated to a time step of 0.1 s. Rise time and fall time of this response were measured as the time from task onset or offset to 50% of the peak response amplitude. This was compared between contrasts using one-tailed Mann-Whitney U-tests with Bonferroni correction (to reduce multiple comparisons, contrasts were not compared between isotropic and linear sequences, or between b200-dfMRI and b1000-dfMRI). *$p < 0.05$, **$p < 0.001$. Cohen's d effect sizes and p-values are shown in Table S4.

| | | Isotropic | | Linear | | |
|---|---|---|---|---|---|---|
| | | b200-dfMRI | b1000-dfMRI | b200-dfMRI | b1000-dfMRI | BOLD-fMRI |
| **Visual Task** | | | | | | |
| Isotropic ADC-fMRI | Rise | 2.23 (0.0030) | n.s. | | | 2.93 (0.0082) |
| | Fall | 2.32 (0.0002) | 1.66 (0.0074) | | | 3.43 (0.0022) |
| Linear ADC-fMRI | Rise | | | n.s. | n.s. | 3.67 (0.0138) |
| | Fall | | | n.s. | n.s. | 2.86 (0.0031) |
| BOLD-fMRI | Rise | n.s. | n.s. | 2.21 (0.0328) | 2.29 (0.0388) | |
| | Fall | 2.03 (0.0198) | 1.97 (0.0175) | 1.71 (0.0483) | 1.71 (0.0231) | |
| **Motor Task** | | | | | | |
| Isotropic ADC-fMRI | Rise | n.s. | n.s. | | | 2.53 (0.0056) |
| | Fall | n.s. | n.s. | | | 1.73 (0.0096) |
| BOLD-fMRI | Rise | 2.60 (0.0057) | 2.25 (0.0078) | | | |
| | Fall | n.s. | 1.70 (0.0119) | | | |

Table S4: Cohen's d effect sizes and Bonferroni-corrected p-values, displayed as d (p), for the comparison of subject-level response time characteristics across contrasts using one-tailed Mann-Whitney U-tests. To reduce multiple comparisons, contrasts were not compared between isotropic and linear sequences, or between b200-dfMRI and b1000-dfMRI

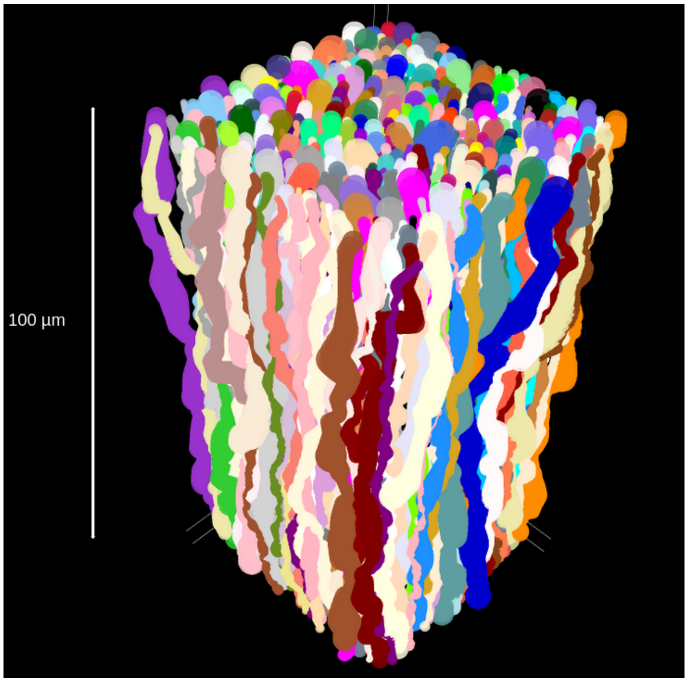


Figure S9: A representative numerical phantom generated for *in silico* experiments using the CATERPillar tool [39]. Axon diameters were drawn from a Gamma distribution with mean of 1 $\mu$m. Axonal beading occurred periodically, with an amplitude equal to 0.3 times the average axon radius. Tortuosity (the ratio of the total axon length to the straight-line distance between its starting and ending points) had an average value of 1.2. The intracellular volume fraction was around 50%.

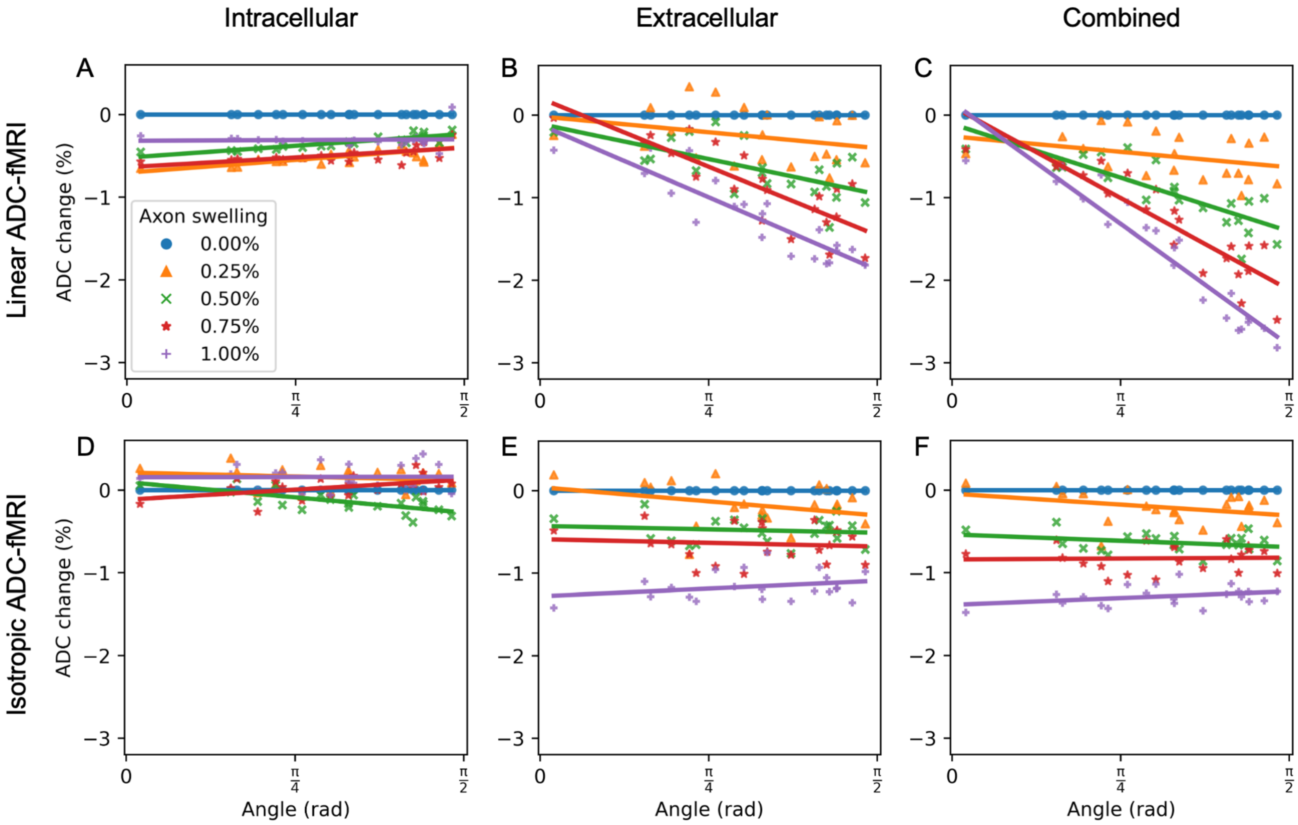


Figure S10: Simulated ADC changes when *in silico* axons swell to 0-1% of their original volume, plotted against fibre angle. This is shown for the intracellular (A & D), extracellular (B & E), and combined (D & F) signal for linear and isotropic ADC-fMRI respectively. Note the combined signal corresponds to that shown in Figure 3 in the main text. Note also that the small decrease in ADC with axonal swelling in the intracellular compartment, particularly for linear encoding, is due to the swelling mechanism implemented in the simulation, where the axon diameter at each position along the axon is swollen by the same percentage. This results in a more beaded axon in the swollen vs rest condition, which is known to reduce diffusivity [91]. Here, the ADC % change in the combined signal is larger than in the extracellular signal due to being measured from a smaller initial ADC (Figure S11) as well as the increase in intracellular contributions to the signal (see Results).

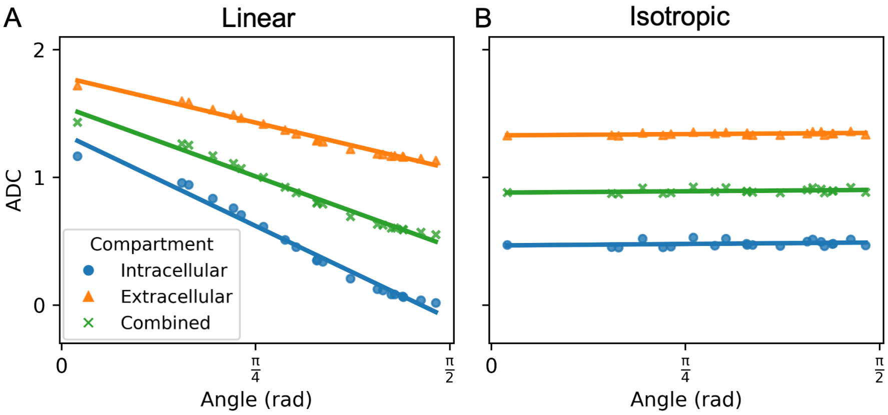


Figure S11: *In silico* ADC measurements at different fibre angles at baseline (i.e. with no axon swelling) for linear ADC-fMRI (A) and isotropic ADC-fMRI (B).

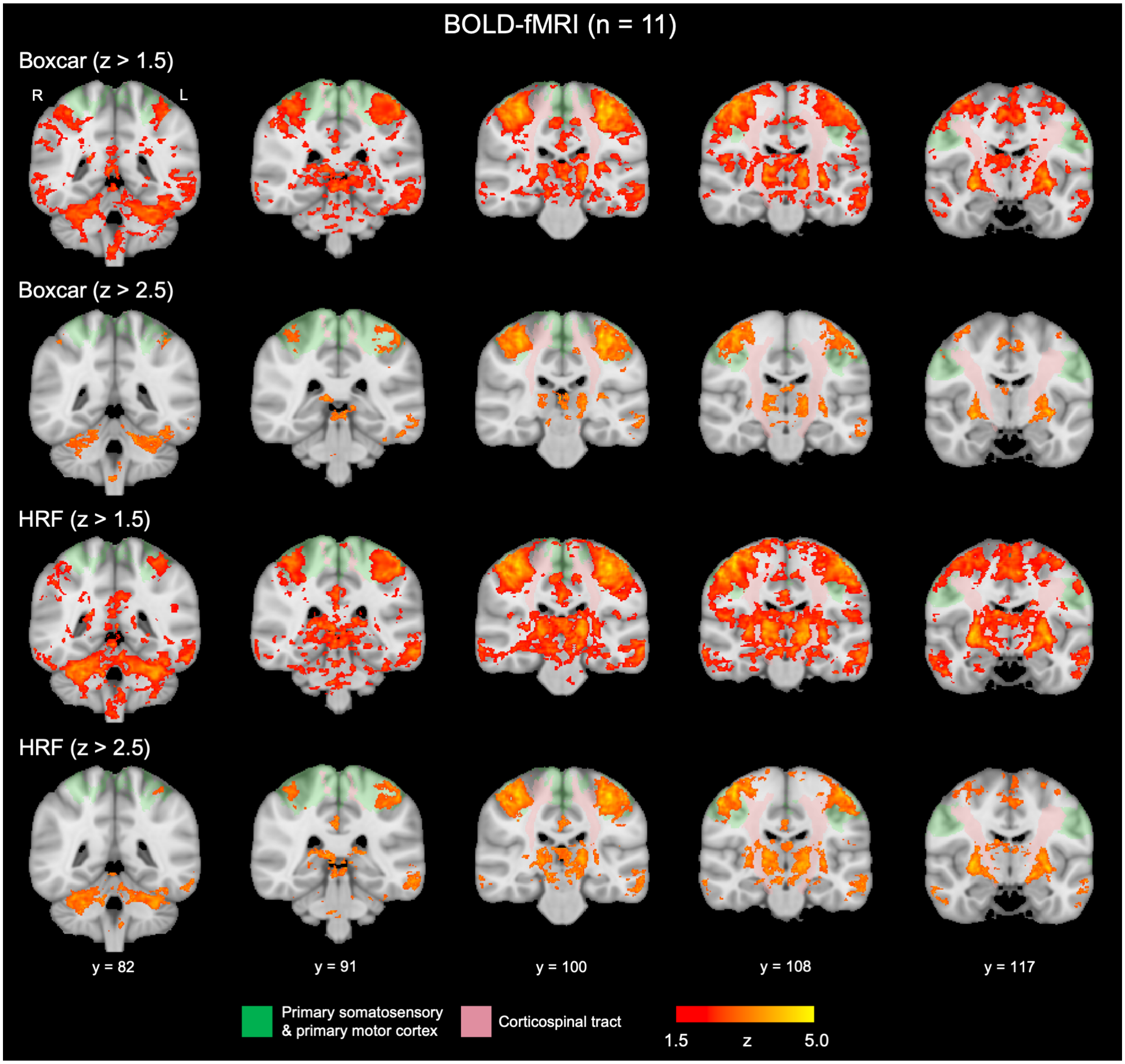


Figure S12: BOLD-fMRI spatial maps showing response to the motor task. The colour bar shows z-score following group-level cluster correction. For comparison with other contrasts, $z \geq 1.5$ ($p < 0.05$) was used for cluster correction; for clarity, results following cluster correction at $z \geq 2.5$ ($p < 0.05$) are also shown here. For both cluster thresholds, spatial maps are shown for the response to the task modelled as a boxcar function and as the convolution of the boxcar function with the canonical haemodynamic response function (HRF). For anatomical reference, Juelich atlas regions defining the corticospinal tract (pink) and the somatosensory and motor cortices combined (green) are overlaid on the MNI152 standard template.

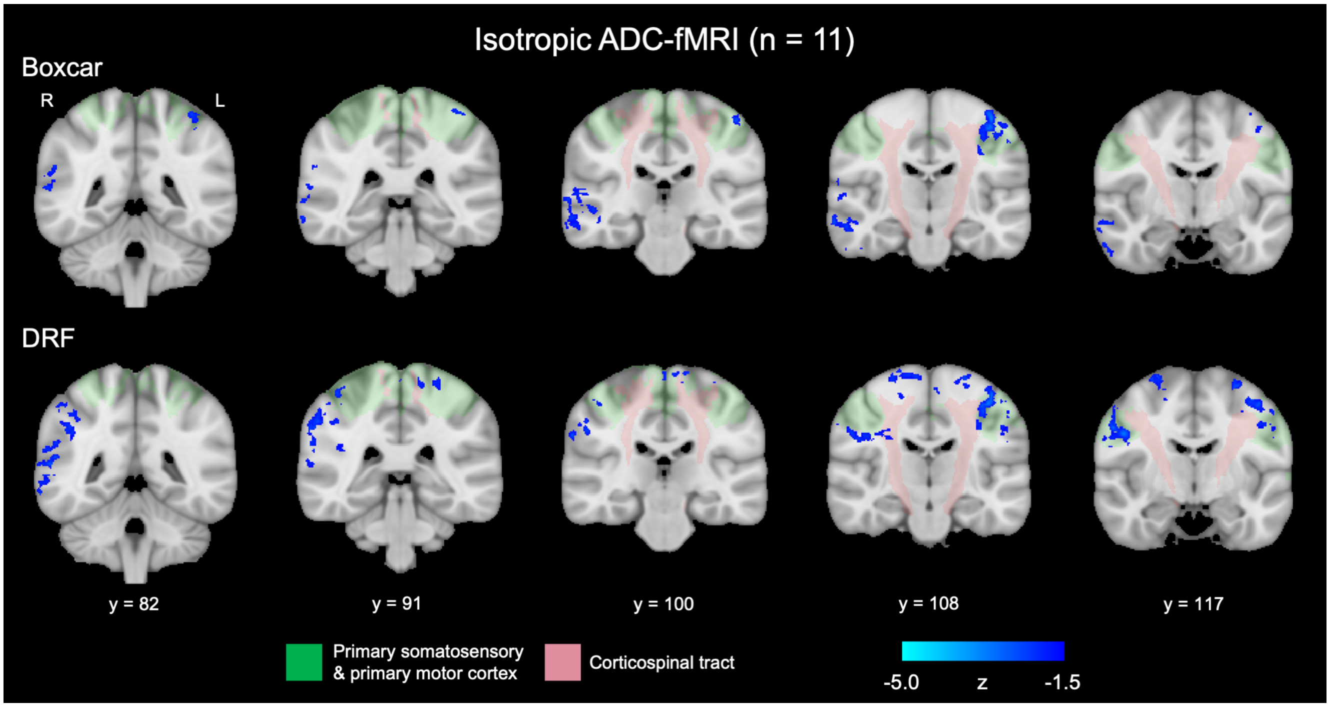


Figure S13: Isotropic ADC-fMRI spatial maps showing response to the motor task with different response functions. The colour bar shows group-level z-score following group-level cluster correction ($z \geq 1.5$, $p < 0.05$). Spatial maps are shown for the response to the task modelled as a boxcar function, and modelled as the convolution of the boxcar function with the diffusion response function [17] (DRF). For anatomical reference, Juelich atlas regions defining the corticospinal tract (pink) and the somatosensory and motor cortices combined (green) are overlaid on the MNI152 standard template.

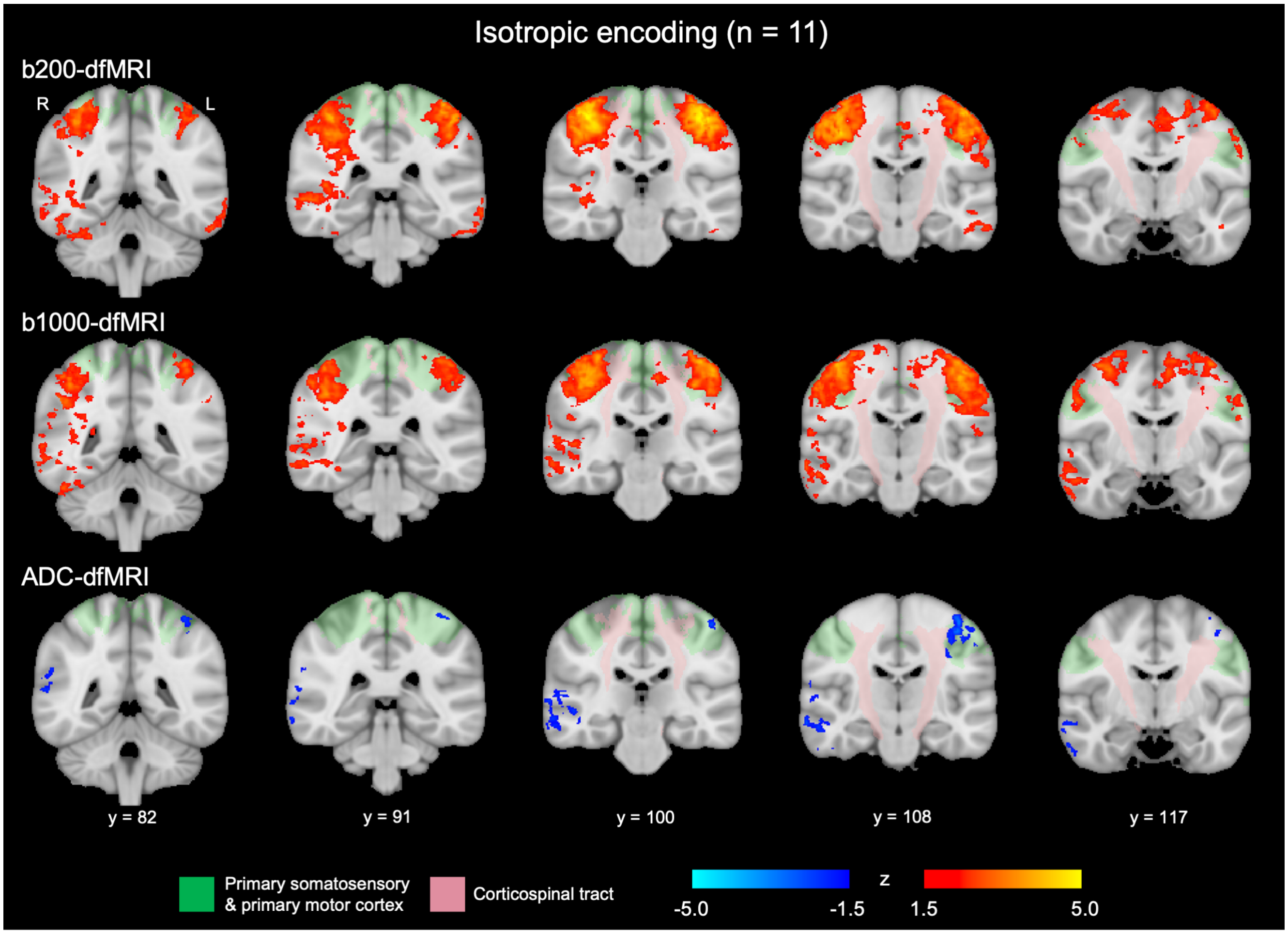


Figure S14: ADC-fMRI, b200-dfMRI and b1000-dfMRI spatial maps for isotropic encoding, showing response to the motor task. The colour bars show z-scores following group-level cluster correction ($z \geq 1.5$, $p < 0.05$). For anatomical reference, Juelich atlas regions defining the corticospinal tract (pink) and the somatosensory and motor cortices combined (green) are overlaid on the MNI152 standard template.

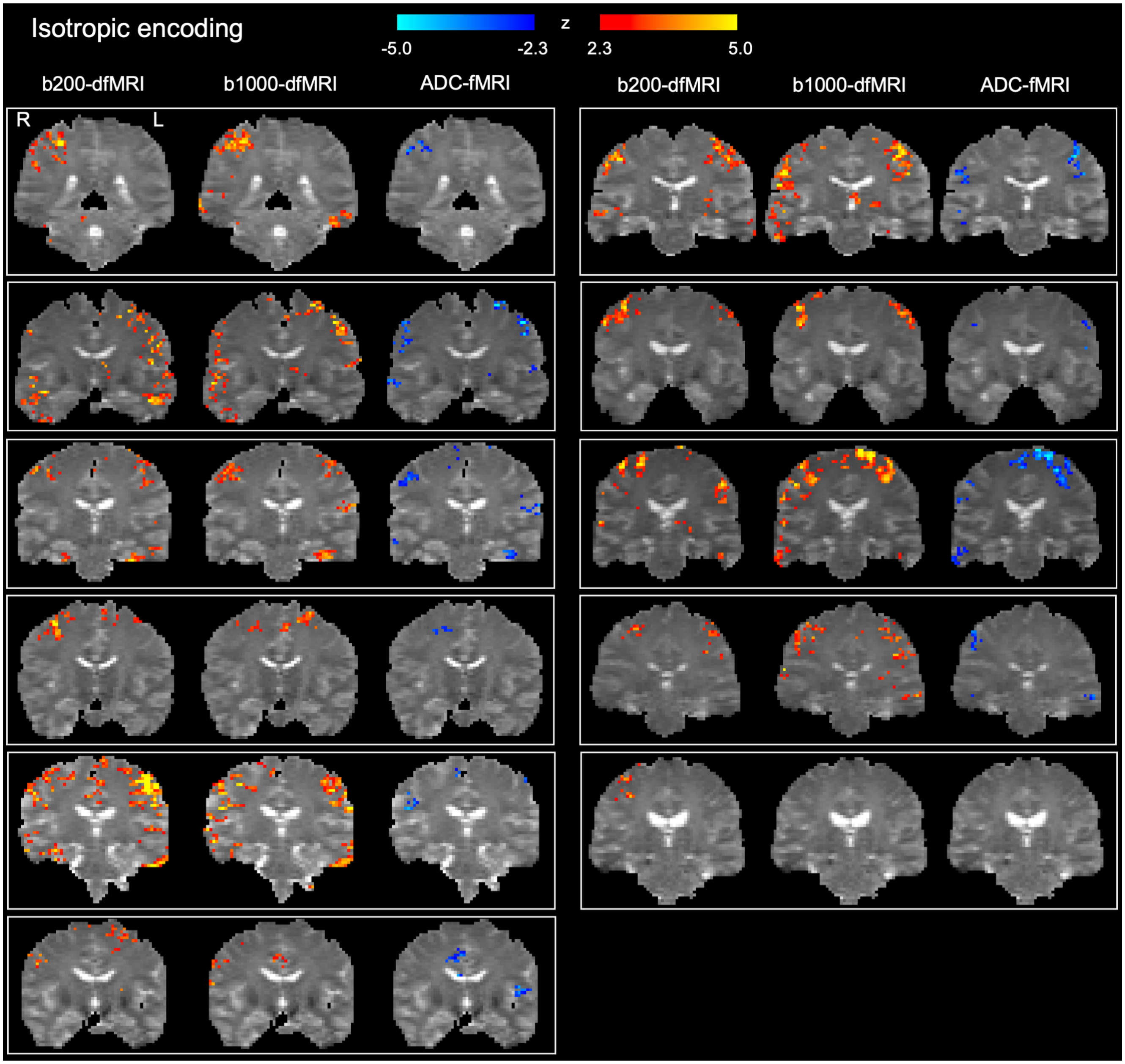


Figure S15: Subject-level motor task spatial maps for isotropic encoding (n = 11). The colour bars show z-scores for b200-dfMRI, b1000-dfMRI and ADC-fMRI following subject-level cluster correction ($z \geq 2.3$, $p < 0.05$).

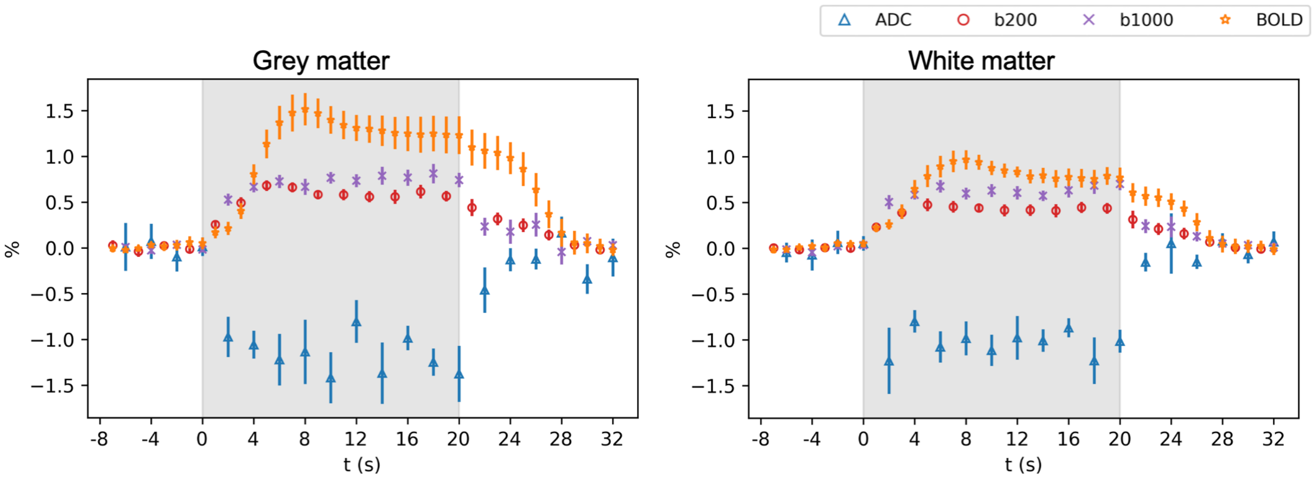


Figure S16: Motor task response in active grey and white matter voxels from subject-level spatial maps. Plots show the average timecourse, with bars indicating the standard error across subjects (n = 11). Subject-specific timecourses were averaged across epochs and across voxels in spatial maps following subject-level cluster correction ($z \geq 2.3$, $p < 0.05$). Time is given in reference to the task onset, with the task stimulation duration indicated by the shaded area.

### High b-value Acquisition and SNR Comparison

To explore the sensitivity of ADC-fMRI at higher b-values, we acquired linear ADC-fMRI datasets (n = 3) with b-values [1000, 2000] s mm$^{-2}$ during visual stimulation. However, we found no task-associated activation in ADC-fMRI timeseries due to the lower SNR of the MRI signal at high b-values. We calculated the image SNR for each b-value timeseries, defined as the mean voxel intensity divided by the standard deviation of the residual noise removed from the voxel during NORDIC denoising (see Methods). Image SNR maps are shown for all ADC-fMRI acquisitions in Figure S17, with mean values in Table S5. For each b-value timeseries and ADC-fMRI timeseries, in addition to BOLD-fMRI, we also calculated temporal SNR as the mean voxel intensity divided by the voxel standard deviation over time, scaled by a factor of $(\mathrm{TR})^{-\frac{1}{2}}$ (Figure S18, Table S6). Both image SNR and temporal SNR are greatly reduced in the high b-value acquisition, which appears to outweigh any increases in sensitivity to diffusion, resulting in no detection of activity with this high b-value ADC-fMRI timeseries.

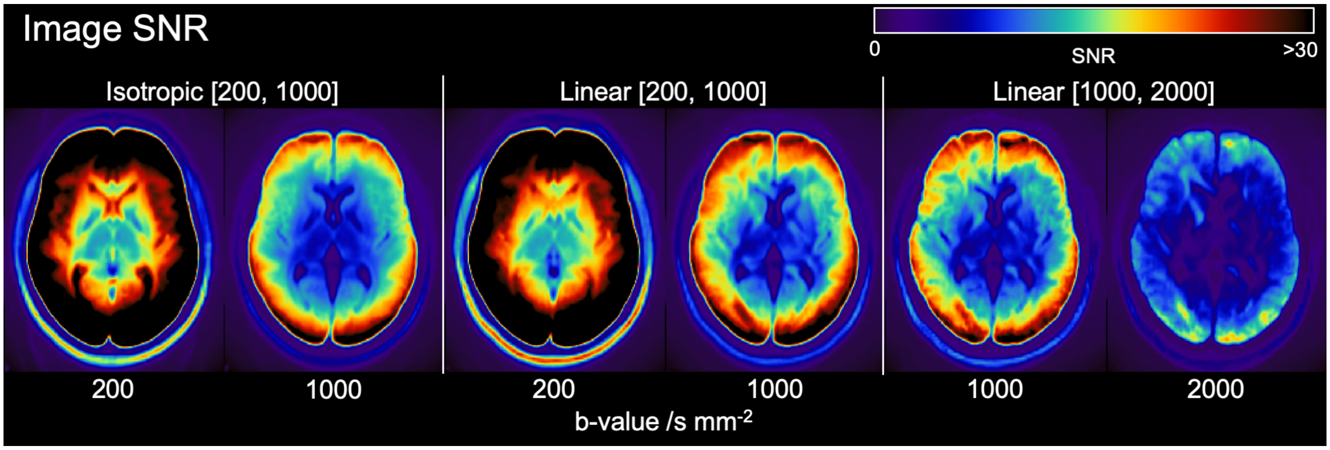


Figure S17: Group-average image SNR for visual task dfMRI acquisitions for isotropic dfMRI with b = 200, 1000 s mm$^{-2}$ (n = 12), linear dfMRI with b = 200, 1000 s mm$^{-2}$ (n = 10), and linear dfMRI with b = 1000, 2000 s mm$^{-2}$ (n = 3).

| Sequence | 200 | 1000 | 2000 |
|---|---|---|---|
| Isotropic [200,1000] | 29.7 (26.7–31.4) | 14.9 (13.4–16.0) | |
| Linear [200,1000] | 30.0 (25.7–32.9) | 16.4 (14.1–18.1) | |
| Linear [1000,2000] | | 14.6 (13.0–15.7) | 8.4 (7.4–8.9) |

Table S5: Image SNR values for visual task dfMRI acquisitions for isotropic dfMRI with b = 200, 1000 s mm$^{-2}$ (n = 12), linear dfMRI with b = 200, 1000 s mm$^{-2}$ (n = 10), and linear dfMRI with b = 1000, 2000 s mm$^{-2}$ (n = 3). Subject-level image SNR was calculated as the average of voxels within the brain. Values show the mean (range) of subject-level image SNR values.

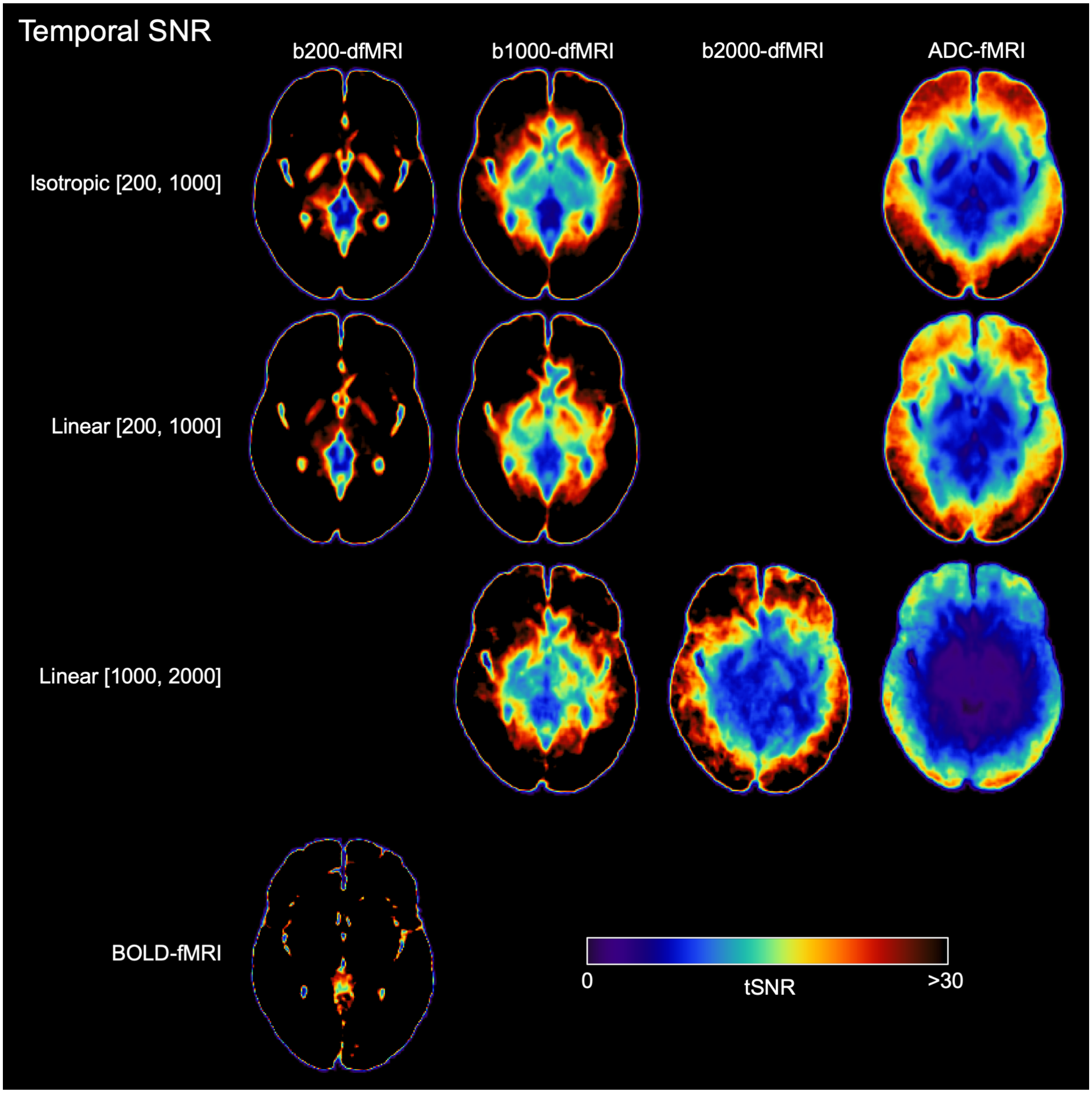


Figure S18: Group-average temporal SNR for visual task data for isotropic dfMRI with b = 200, 1000 s mm$^{-2}$ (n = 12), linear dfMRI with b = 200, 1000 s mm$^{-2}$ (n = 10), linear dfMRI with b = 1000, 2000 s mm$^{-2}$ (n = 3), and BOLD-fMRI (n = 7).

| Sequence | b200-dfMRI | b1000-dfMRI | b2000-dfMRI | ADC-fMRI |
|---|---|---|---|---|
| Isotropic [200,1000] | 64.5 (54.5–78.2) | 42.1 (36.1–49.3) | | 27.0 (22.5–32.2) |
| Linear [200,1000] | 69.0 (62.1–76.0) | 46.0 (38.5–54.5) | | 24.9 (21.9–28.0) |
| Linear [1000,2000] | | 44.2 (37.3–48.0) | 29.4 (26.1–32.8) | 15.2 (14.1–16.6) |
| BOLD-fMRI | 85.7 (76.1–89.3) | | | |

Table S6: Temporal SNR values for visual task data for isotropic dfMRI with b = 200, 1000 s $mm^{-2}$ (n = 12), linear dfMRI with b = 200, 1000 s $mm^{-2}$ (n = 10), linear dfMRI with b = 1000, 2000 s $mm^{-2}$ (n = 3), and BOLD-fMRI (n = 7). Subject-level temporal SNR was calculated as the average of voxels within the brain. Values show the mean (range) of subject-level temporal SNR values.

### Model-Free Frequency Analysis

In order to assess whether BOLD-fMRI could detect signals time-locked to the visual stimulation, with no assumptions about the response characteristics, we performed model-free analysis of the power spectrum of the BOLD-fMRI signal in the optic radiation, following the method of Schilling et al. [37].

#### Methods

BOLD-fMRI data for each subject from the visual task (following all preprocessing and temporal filtering) were transformed to MNI space and resampled to 2 mm isotropic resolution, then each voxel timeseries was temporally normalised and converted to a percentage signal change. We used the optic radiation mask from the Juelich atlas, masked to eliminate grey matter voxels. In order to assess the white matter BOLD signal with minimal contributions from partial volume with grey matter, we also eroded this mask with both a 3 x 3 x 3 $mm^3$ kernel and a 5 x 5 x 5 $mm^3$ kernel (Figure S19A). These three optic radiation masks were resampled to match the resolution of the functional data.

The average BOLD signal within each region of interest (ROI) was extracted and averaged across subjects (Figure S19). The power spectral density of this group-averaged signal was estimated using the periodogram method, and the power at the task frequency (0.0333 Hz; 1/30 s) was measured. To measure the significance of this power, we generated a null distribution using bootstrap analysis [37], as follows. We sampled 10,000 bootstrap samples from volumes of 30 x 30 x 22 $mm^3$, taking the average signal across subjects within each sampled volume then measuring the power at the task frequency in the sampled timeseries. We then calculated a p-value for the original data by comparison with this null distribution. In the original work [37], the authors generated the null distribution from resting-state data. As we do not have resting-state data with acquisition parameters matching those of the task experiments, we derived a null distribution by bootstrapping from the visual task data, but only including patches from the anterior half of the imaging volume (i.e. with no overlap with primary visual processing regions or optic radiation).

#### Results

In all three optic radiation ROIs, there was significant power at the task frequency ($p < 0.05$) compared to bootstrap samples from non-visual processing areas of the brain in the same data, demonstrating that the BOLD-fMRI signal can capture activity, at an ROI level, which is time-locked with the task. The more eroded optic radiation masks, which are more specific to deep white matter, exhibit a different response to the stimulus, with a slower rise (as previously shown [7]). This altered response is likely the reason for the lack of detection of deep white matter voxels in the voxel-wise general linear model analysis in the main text. These results exemplify the heterogeneity of the haemodynamic response across the brain which obstructs BOLD-fMRI analysis of simultaneous grey and white matter activity (for example general linear model analysis or resting-state functional connectivity).

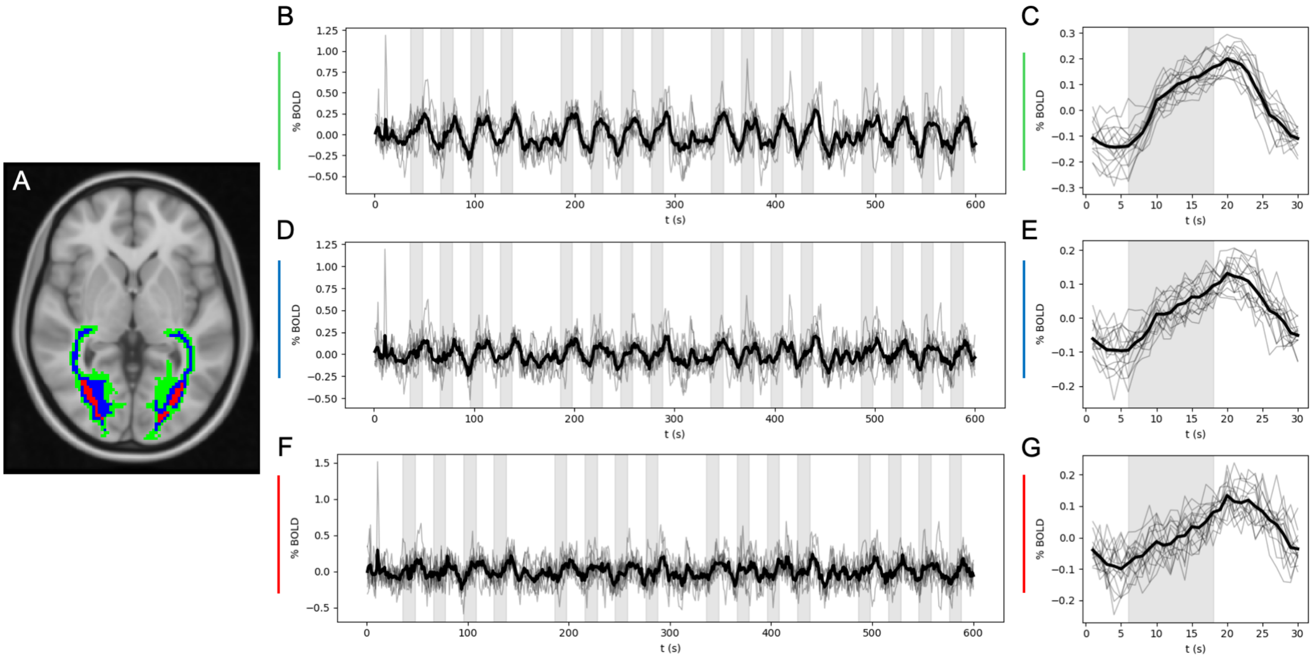


Figure S19: ROI-average BOLD-fMRI signal during visual stimulation. A) Three optic radiation masks were used; the original Juelich atlas region was masked to only include white matter voxels (green), which was then eroded with a 3 mm kernel (blue) and a 5 mm kernel (red) and resampled to 2 mm resolution. B, D, F) Within each of these optic radiation ROIs, the mean BOLD-fMRI signal was measured from each subject (grey lines) and averaged across subjects (black lines). C, E, G) From the group-averaged BOLD-fMRI timeseries, each task epoch is plotted (grey lines) and averaged across epochs (black lines). Plots B & C correspond to the full optic radiation, D & E correspond to the 3 mm-eroded mask, and F & G correspond to the 5 mm-eroded mask. Stimulus blocks are indicated by the shaded area.
